# Interdependence of primary and secondary somatosensory cortices for plasticity and texture discrimination learning

**DOI:** 10.1101/2023.04.25.538217

**Authors:** Anurag Pandey, Sungmin Kang, Nicole Pacchiarini, Hanna Wyszynska, Aneesha Grewal, Alex Griffiths, Imogen Healy-Millett, Zena Masseri, Neil Hardingham, Joseph O’Neill, Robert C. Honey, Kevin Fox

**Author notes:** Address for correspondence: Prof Kevin Fox, School of Biosciences, Cardiff University, Museum Avenue, CF10 3AX.

## Abstract

Feedforward and feedback pathways are important for transfer and integration of information between sensory cortical areas. Here we find that two closely connected cortical areas, the primary (S1) and secondary somatosensory cortices (S2) are both required for mice to learn a whisker-dependent texture discrimination. Increased inhibition in either area (using excitatory DREADDs expressed in inhibitory interneurones) prevents learning. We find that learning the discrimination produces structural plasticity of dendritic spines on layer 2/3 pyramidal neurones in vibrissae S1 that is restricted to the basal dendrites and leaves dendritic spines on apical dendrites unchanged. As S2 projects to the apical dendrites of S1 neurones, we tested whether S2 affects LTP-induction in S1. We found that feedback projections from S2 to S1 gates LTP on feedforward pathways within S1. These studies therefore demonstrate the interdependence of S1 and S2 for learning and plasticity in S1.

**HIGHLIGHTS:**

- Both primary (S1) and secondary (S2) somatosensory cortices are necessary for whisker based texture discrimination learning
- S2 feedback connections to S1 gate LTP at feedforward pathways in S1
- S1 undergoes structural plasticity of pre-existing spines during learning
- S1 learning induced plasticity and LTP occurs on basal but not apical dendrites

## Introduction

Cortical areas communicate via a series of feedforward and feedback loops that integrate higher order representations with sensory information conveyed from the periphery via the thalamus (Van Essen et al., 1992). Feedback connectivity is likely to be required to transmit task-specific contextual information to primary cortex to improve perception when feedforward information is noisy, occluded or ambiguous (Bondy et al., 2018; Dura-Bernal et al., 2012; Ebrahimi et al., 2022; Gilbert and Li, 2013; Lamme, 1995; Zipser et al., 1996).

Feedback connectivity to primary cortex may also amplify or tag novel sensory features during learning. Indeed, feedback connections to the cortex are thought to be important for long-term memory consolidation. This is because the hippocampus tends to store recent memories while the neocortex is responsible for storing memory over a longer time period, implying that information is fed back to the neocortex from the hippocampus (Goto et al., 2021; Sawangjit et al., 2018; Squire and Alvarez, 1995). In concert with this idea, feedback connections from perirhinal cortex to barrel cortex and from secondary motor cortex to barrel cortex have both been found to be important for learning (Doron et al., 2020; Miyamoto et al., 2016). Therefore, the idea that learning and memory operate by inducing synaptic plasticity (Goto et al., 2021; Martin et al., 2000) leads to the conclusion that feedback connections should affect synaptic plasticity in the cortex during learning, a point that we test in this study.

Feedback connections from higher order cortical areas typically terminate strongly on apical dendrites of pyramidal neurones in layer 1 of the cortex (Cauller, 1995; Schuman et al., 2021). In primary somatosensory cortex, higher order somatosensory areas convey information via the apical dendrites (Mao, Kusefoglu et al. 2011, Minamisawa, Kwon et al. 2018) while the basal dendrites carry information, either directly or indirectly, from ascending thalamic pathways (Hooks et al., 2011). Feedforward and feedback information therefore converge onto single neurones at different dendritic locations.

We recently found that the degree of plasticity shown by the synapses located on apical and basal dendrites during sensory deprivation differ markedly. Layer 2/3 pyramidal and layer 5 intrinsic bursting neurones exhibit synaptic plasticity on their basal rather than their apical dendrites (Pandey et al., 2022; Seaton et al., 2020). Basal dendrite synapses undergo plasticity comprising an increase in the formation and survival of newly formed synapses and enlargement of pre-existing synapses. Both forms of structural plasticity are dependent on αCaMKII-autophosphorylation (Seaton, Hodges et al. 2020), which is in turn known to be important for LTP and experience-dependent potentiation (Giese, Fedorov et al. 1998, Hardingham, Glazewski et al. 2003). However, the apical dendrites do not exhibit either form of structural plasticity during sensory deprivation (Seaton et al 2020).

It is not known whether this dichotomy in synaptic behaviour is an oddity of sensory deprivation or a more general phenomenon. We therefore investigated the mechanisms by which synaptic plasticity might be induced on different dendritic loci in primary somatosensory cortex during natural learning. We made use of a texture discrimination behaviour that requires the animal to forage in the dark for a food reward hidden in one of two bowls, distinguishable only by their outer texture. This behavioural assay is modality specific and relies on the whiskers for performance (Pacchiarini et al., 2020). To investigate the role of cortical feedback in this process, we focused on the feedback projections from the secondary somatosensory cortex (S2) to the primary somatosensory cortex (S1) (Minamisawa et al., 2018). Specifically, we studied the vibrissae representation in S1 (S1) and S2 (vS2). We initially tested whether learning requires S1 and S2 and in particular the whisker receptive areas, before investigating structural plasticity and LTP induction mechanisms in S1, and its dependence on S2. Our findings reveal that both S1 and S2 are necessary for learning the texture discrimination and that learning produces structural plasticity on the basal dendrites of layer 2/3 neurones in S1 not the apical dendrites; we further show that a novel feedback mechanism exists, by which S2 gates plasticity on the basal dendrites of S1 cortical neurones.

## Results

### 1. Texture discrimination learning

We adapted a rodent discrimination assay (Birrell and Brown, 2000) to test whether mice could learn a tactile texture discrimination. The method involves the mouse discriminating between different textures present on the outer surface of a bowl. The bowls are filled with sawdust and one of the textures signals a reward (a cocoa-pop) that is hidden in the sawdust (Figure 1A, Supplementary Figure 1). The mouse must dig to find the reward, which affords an easy method of scoring the behaviour. The mouse is allowed to explore either of the two chambers containing the bowls until it starts to dig in one of the bowls, whereupon access to the other chamber is prevented by the closing of a door. The behaviour takes place in dim red light, with the odour of the cocoa-pop reward masked. Previous studies have shown that mice require their whiskers to learn the texture discrimination (Pacchiarini et al., 2020). In early experiments, we ran 20 trials per day for two days, but in later experiments we increased the number to 24 trials per day for 3 days. We found that mice learned the discrimination quickly (Figure 1B), improving from a mean of 50% correct to a mean of 70% correct within two days, confirming previous findings (Pacchiarini et al., 2020).

**Figure 1.**
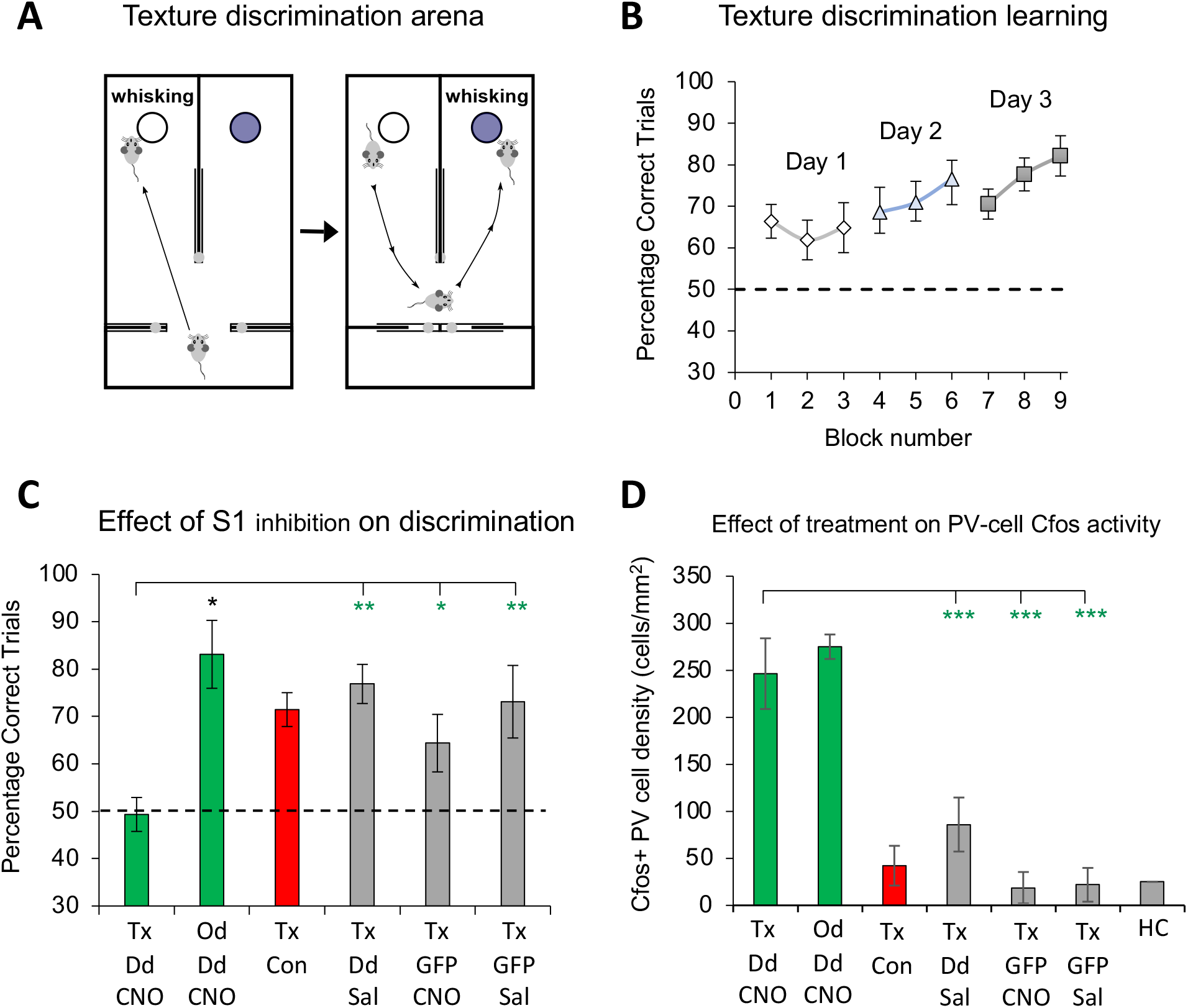
Learning performance in a texture discrimination assay with and without inhibiting S1 barrel cortex. **A:** The testing arena: Two bowls either side of a partition, only one of which is baited, can be investigated until a choice is made by digging in the sawdust of one bowl. **B:** The percentage of correct choices between the two textured bowls increases over three consecutive days of training. **C:** The percentage of correct trials across different conditions averaged across all days (see Results). From left to right, inhibiting S1 results in chance levels of performance (50%) on a texture discrimination (Tx, Dd, CNO; texture DREADD, CNO), but has no effect on an odour discrimination (83% correct, green bar, Od, Dd, CNO). Grey bars show three control conditions Texture+DREADD+Saline (Tx, Dd, Sal), Texture+GFP+CNO (Tx, GFP, CNO), Texture+GFP+Saline (Tx, GFP, Sal) (4 mice per group) and the red bar their average. All three conditions are significantly different from the Texture+CNO+DREADD condition. **D:** Cfos positive cells density is shown for the same conditions as shown in **C** plus one extra condition (HC, home caged) where mice had no interaction with the test arena. Cfos+ density is significantly higher only when CNO and DREADD expression are combined (green asterisks denote significance for between group comparison, *p<0.05,**p<0.005, ***p<0.002, black asterisk denotes significant difference from chance levels (50%), *p<0.03).

### 2. Texture discrimination learning depends on barrel cortex

To test whether the texture discrimination depended on normal neuronal activity in the barrel cortex, we inhibited the barrel cortex during the discrimination assay. We injected AAV virus coding for floxed hM3D(Gq) DREADD into the barrel cortex bilaterally in a PV-cre line of mice, aiming our injections at layer 4 of the D2, D6 and B2 barrel-columns (see Methods). The excitatory DREADD was expressed only in PV cells. Approximately 30 minutes before discrimination training began for each mouse, we administered CNO i.p. at a dose of 3.5mg/kg. To control for the effects of transgenic expression in PV cells and any general effects of CNO, we used an experimental design involving DREADD or GFP expression and CNO or saline injection.

A one-way ANOVA conducted on the results from the four treatment groups performing a texture discrimination confirmed that there was a main effect of treatment (F_(3,19)_ = 6.904, p < 0.005, eta-squared = 0.564). Least Significant Difference testing revealed that the scores for each of the three control groups (DREADD+Saline, GFP+CNO, GFP+Saline; grey bars Figure 1C) differed from those in the DREADD+CNO group (green bar; p < 0.05 for the GFP+CNO group and p < 0.005 for the other two control groups). There were no significant differences between the three control groups, for which the red bar (Figure 1C) provides the mean. Furthermore, while the scores for the mice in the control groups differed significantly from 50% correct, (t_(11)_ = 6.014, p < 0.001, Cohen’s d = 1.736), those from the DREADD+CNO group did not, (t_(7)_ = -0.189, p > 0.85, Cohen’s d = -0.067).

We tested the possibility that inhibiting the barrel cortex activity had a non-specific effect that generally prevented learning. To do this, we assessed the effect of the same treatment on mice learning an odour discrimination, where the odours in two bowls indicated whether the reward was present or not, noting that this odour discrimination is not affected by whisker trimming (Pacchiarini et al., 2020). In contrast to the texture discrimination assay, we found that the scores for the mice in the hM3D(Gq)-CNO group, given an odour discrimination (Figure 1C, labelled ‘OD Dd Tx’), differed significantly from 50% correct, t(3) = 4.619, p < 0.03, Cohen’s d = 2.309. We therefore conclude that the barrel cortex is required for learning a texture discrimination and that the manipulation that we used is modality specific.

### 3. Verification and quantification of the DREADD effect

The location of DREADD expression was assessed from post-mortem histology using the native mCherry signal from the DREADD virus and VGlut2 antibodies to stain thalamic axons in the barrel field (Supplementary Figure 2). Expression of DREADDs in PV cells covered an average area of 500µm^2^ in each hemisphere (range 185-1090µm^2^). The area of expression was similar in all treatment groups, but appeared slightly larger in the odour discrimination group (mean - 683 µm^2^ versus 466 µm^2^), though a one-way ANOVA showed that this was not statistically significant (F_(4,41)_=2.32, p=0.075).

The effectiveness of DREADD expression in PV cells was assessed in several ways; first, by measuring cfos expression in PV and non-PV cells inside and outside the area of DREADD expression; second, by single unit recordings in the middle of the expression area; and third, by Neuropixels electrode recordings traversing vibrissae-receptive areas in S1 and S2.

#### CFOS

Cfos immuno-staining showed that PV cells significantly increased their cfos expression as a result of CNO mediated DREADD activation (Figure 1D, Supplemental Figure 2). Cell counts of PV cells expressing cfos above a standard threshold level (see Methods) increased ten fold following CNO activation (Figure 1D). A two-way ANOVA for DREADD (DREADD vs GFP) and CNO (CNO vs Saline) revealed an interaction between DREADD and CNO (F_(1,1)_=5.86, p=0.0296). Post hoc t-tests showed this was due to the DREADD expressing mice treated with CNO being different from all other treatment combinations (p<0.002, Figure 1D). Conversely non-PV cells decreased their cfos expression levels significantly compared to controls within the same regions of interest in the same sections (reduction to 79% of control levels, F_(1,9)_=6.89, p=0.0303, one-way ANOVA). A similar result was found when intensity levels were analysed rather than cell counts; PV cells increased their cfos intensity levels by more than 3 fold and non-PV cells showed levels reduced to 69% of control.

#### SINGLE ELECTRODE RECORDINGS

Single cell recordings using carbon fibre micro electrodes showed that a single dose of 3.5mg/kg of CNO was sufficient to reduce responses noticeably within 5-10 minutes and abolish responses to the principal whisker in cells located in layer 2/3 of the barrel cortex within 15 minutes (Figure 2A). A CNO dose of 1.75mg/kg reduced the principal whisker responses to 36± 6% of control baseline values but did not completely abolish responses over a period of 2 hours. Injection of CNO in animals lacking DREADDs had no detectable effect on the principal whisker responses (Figure 2A). Both CNO doses produced statistically significantly reduction in responses compared to CNO alone (3.5mg/kg dose, F_(1,12)_=22.7, p<0.001; 1.75mg/kg dose F_(1,11)_=67.4, p<0.0001) or to their own baseline control levels (ANOVA: 3.5mg/kg dose, F_(1,11)_=15.5, p<0.003; 1.75mg/kg dose F_(1,9)_=91.9, p<0.0001; PAIRED T-TEST: 3.5mg/kg dose, t_(2)_=25.3, p<0.005; 1.75mg/kg dose t_(4)_=9.4, p<0.001).

**Figure 2.**
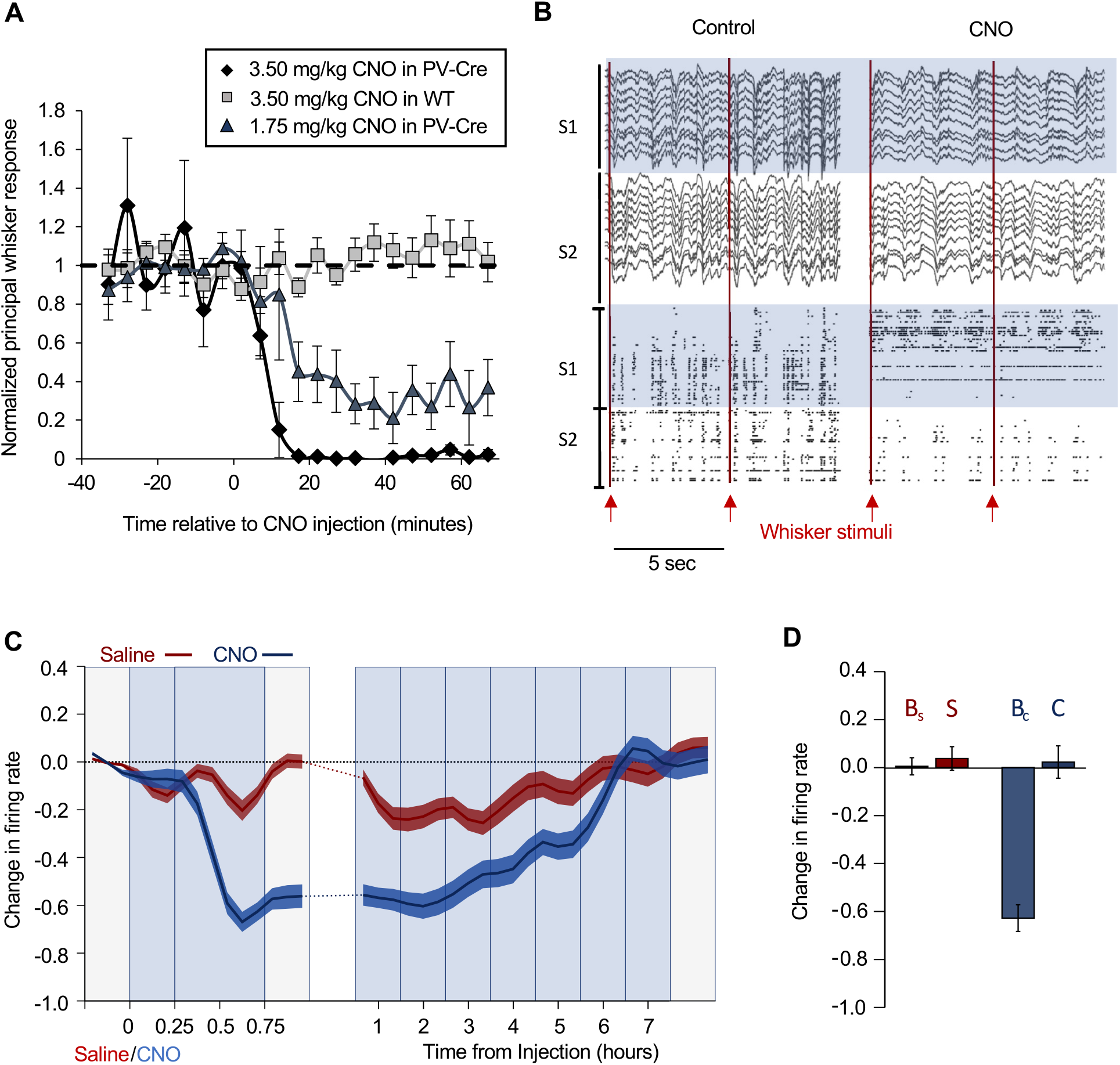
Electrophysiological quantification of DREADD effect. **A:** Inhibition of responses to principal whisker stimulation using two concentrations of CNO (dark grey triangles and black diamonds) in mice expressing hM3Dq in PV cells compared the higher CNO concentration in animals lacking hM3Dq expression (grey squares). **B:** Neuropixels recording showing local field potentials and spike rasters for the same spikes recorded before and after injection of CNO in a mouse expressing hM3Dq DREADD in PV cells in S1 barrel cortex. Red lines show timing of C2 whisker stimulation. Top: S1 (blue) shows reduction in delta power while S2 is moderately affected. Continuously firing cells after CNO are presumed interneurones. Bottom: S1 shows complete loss of sensory responses and network activity (blue). S2 shows reduction in sensory responses and network activity. **C:** Time course of action of a single CNO injection on firing rate in 48 neurones located in S1 recorded with a Neuropixels probe in a freely moving rat expressing hM3Dq in inhibitory neurones. Light grey background represents periods of exploration and blue background periods of rest. The blue line shows the effect of CNO activating the DREADDs and the red line a saline injection for the same cells on different days. The change in firing rate (Instantaneous rate-baseline)/(Average rate+baseline) is plotted together with SEM. Note the change in time-base for the period just after CNO injection compared with recovery. **D:** Difference histograms for deviations from baseline. Bs baseline before saline injection. S average during saline injection period. Bc baseline before CNO injection. C average during CNO injection relative to baseline. NB: all averages are of activity during rest (i.e. excluding exploration periods).

#### MULTISITE ELECTRODE RECORDINGS

Neuropixels recordings allowed us to view simultaneously both the timing of CNO mediated DREADD action and the area affected. We made recordings from 3.5mm deep electrode penetrations angled to run through S1 barrel cortex and the whisker responsive area of S2 (see Methods). Recordings from 350 electrodes per penetration yielded approximately 200 cells per animal following a cell clustering analysis (See Methods). In each experiment, we chose the whisker that drove the best aggregate responses in S1 and S2 simultaneously.

Network activity was profoundly disrupted within 5 minutes of CNO injection in experiments where DREADDs had been injected in S1. While in control conditions we saw synchronous delta-wave activity and associated bursts of action potentials in control conditions, following CNO injection we found that delta-wave power and the associated spike activity were reduced (Figure 2B, Supplementary Figure 3). Simultaneously, many action potentials that were not previously observed, began firing tonically at an average firing rate of 10-20Hz, while the delta-wave associated bursts of action potentials ceased in S1, but continued in S2 (Figure 2B, Supplementary Figure 3). The local field potentials followed a similar trend, showing a large loss of power in the delta-wave band (0.5-4Hz) in S1 (82% decrease, p<0.001) and a smaller decrease in S2 (21% decrease, p<0.001) (Supplementary Figure 3). These changes could not be replicated by injecting CNO into a mouse lacking DREADD expression (Supplemental Figure 3D).

Sensory responses were affected in a variety of ways (Supplementary Figure 4), with cells either decreasing or losing their sensory response all together, or in a small number of cases, with cells increasing their response. Some cells, presumed inhibitory neurones from their spike and firing properties (see Methods), increased their firing rate to reveal an inhibitory response during stimulation (Supplementary Figure 4). Quantification of sensory responses to single whisker stimulation (usually the C2 whisker) showed responses were strongly decreased in S1 (for 45.0% of excitatory cells and 48% of inhibitory cells) and decreased slightly for a small percentage of S2 cells (12.2% of excitatory cells and 7.7% of inhibitory cells) following DREADD activation (Supplementary Figures 4, 5A, B). The loss of evoked field potentials in S1 mirrored the loss of spike activity (Figure 2B). In conclusion, activation of excitatory DREADD in inhibitory cells in S1 profoundly inhibited network activity and reduced sensory responses for approximately 46% of S1 neurones, with small effects on S2.

Finally, we estimated the duration of the action of a single injection of CNO by recording from an awake rat chronically implanted with a Neuropixels probe in S1. We found that induction of the effect was rapid (less than 15 minutes after intraperitoneal administration of CNO), and that the network activity of S1 in awake and resting states returned to baseline within 6 hours (Figure 2C). Since the texture discrimination sessions lasted approximately 1.5 hours, and CNO was administered 0.5 hours before training began, excitatory neurone activity was inhibited for approximately 4 hours beyond texture discrimination learning.

### 4. Texture discrimination learning causes structural plasticity in the barrel cortex

We tested whether layer 2/3 neurones underwent structural plasticity in the barrel cortex as a consequence of learning the texture discrimination. Sparse labelling with fluophore was achieved by injecting floxed GFP mixed with a limiting titre of an AAV virus expressing Cre protein under the control of an αCaMKII promoter (see Methods). Previous studies indicated that spines on apical and basal dendrites exhibited different plasticity (Pandey et al., 2022; Seaton et al., 2020), therefore we treated spine location as a factor in the analysis. Spine imaging was conducted for two baseline periods, a period immediately after the third day of texture discrimination and finally, after a further three days in the home cage without any intervening texture discrimination trials (see Methods).

In a control group that only experienced their home cage, we found that baseline spine formation and elimination were approximately in equilibrium so that no net gain or loss of spines occurred from one imaging session to the next (Figure 3A). Similarly, spine formation and elimination were in equilibrium during the baseline period before the texture discrimination.

**Figure 3.**
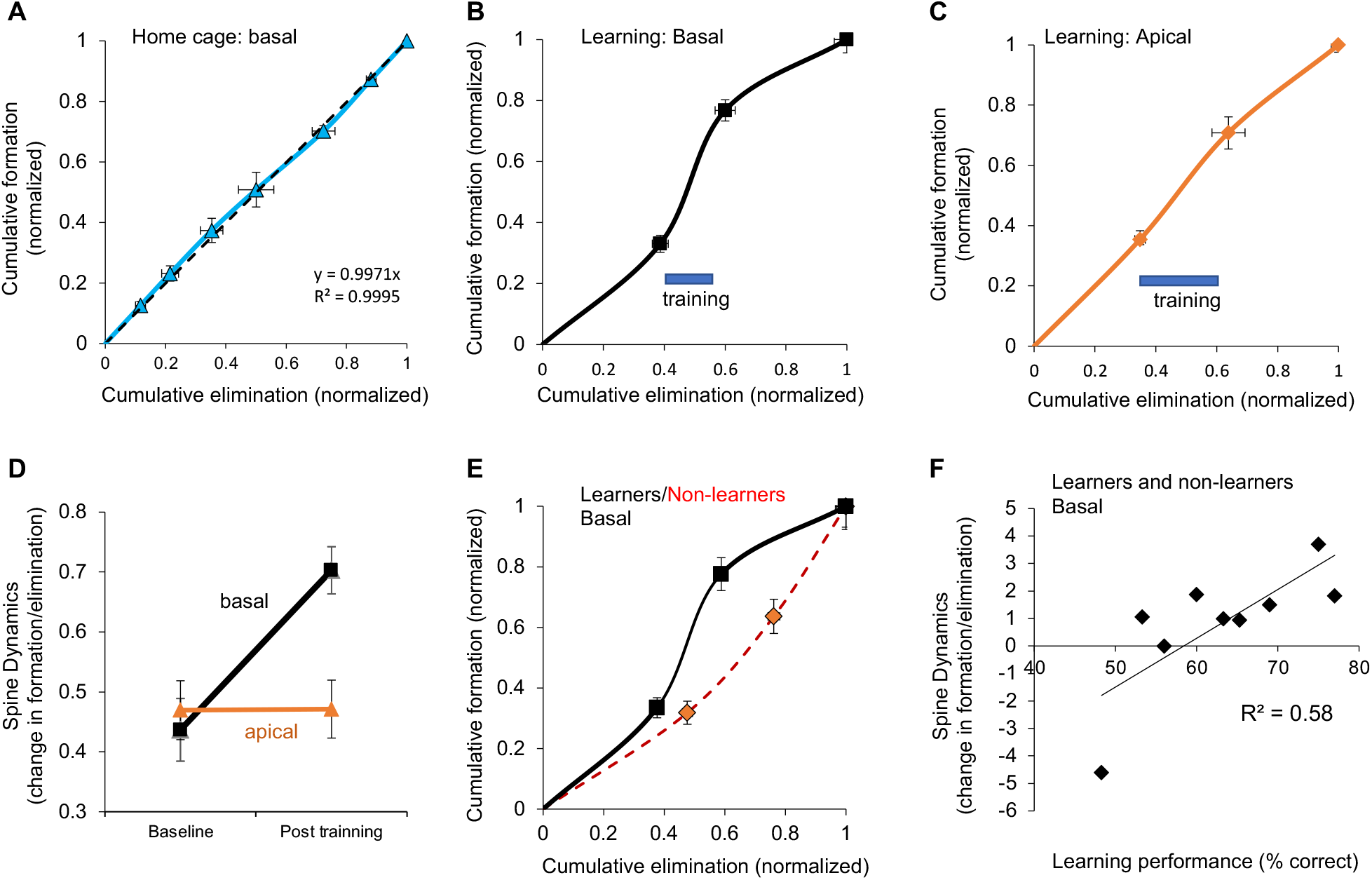
Spine dynamics during texture discrimination learning. **A:** Spine formation and elimination are in equilibrium for basal dendrites of home cage mice. **B:** Spine equilibrium is broken during texture discrimination training on basal but not (**C**) apical dendrites. Note that the origin corresponds to B1, the second point to B2, the third to T1 and the last point to F1. **D:** Spine equilibrium index for apical (orange) and basal dendrites (black). **E:** Spine dynamics for mice that learned (black line) plotted separately from those that did not (non-learners, red dashed line). **F:** Correlation between learning performance and spine dynamics plotted for each animal.

However, following three days of texture discrimination, the balance between formation and elimination changed significantly for spines located on basal dendrites (Figure 3B,D) but not apical dendrites (Figure 3C,D). A two-way ANOVA for learning and spine location showed an effect of training (F_(2,2)_ = 6.75, p<0.002) and an interaction between training and spine location (F_(2,2)_=7.49, p<0.001). For basal dendrites, the rate of spine elimination was reduced to 45% of baseline levels during learning (F_(2,44)_ = 10.03, p<0.0003) while the rate of formation was not changed significantly (F_(2,44)_=2.11, p=0.133)). For apical dendrites, no changes were seen either for elimination rates (F_(2,38)_ =1.86, p=0.17) or formation rates (F_(2,38)_ =0.58, p=0.57).

#### CORRELATION WITH LEARNING

In a pilot study we found that mice fitted with a head plate for S1-imaging were able to learn the texture discrimination within two days (average 76% correct trials, n=2). However, when we repeated the study in combination with spine imaging, we found some variability in performance between mice. Using a binomial test to gauge performance, 3/9 mice did not learn at the α=0.05 significance level within three days (72 trials). We therefore assessed if there were any differences in spine dynamics in mice that learned versus those that did not learn (Figure 3E). A three-Way ANOVA, looking at the effect of spine location, learning outcome and time-point (baseline, texture discrimination and probe trial periods), showed a main effect of time-point (F_(1,1)_=6.0, p<0.02) and an interaction between learning outcome and spine location (F_(1,1)_=5.2, p<0.03). Separate two-way ANOVAs for the mice that learned and those that did not, showed a main effect of time (F_(1,1)_=7.4, p<0.01), spine location (F_(1,1)_=9.5, p<0.005) and an interaction between the two for mice that learned (F_(1,1)_=5.8, p<0.03) but no significant effects at all for those that did not (F_(3,25)_=0.54, p=0.65). In further corroboration, we found a significant correlation between learning outcome and basal dendrite plasticity (R^2^=0.58, F_(1,7)_=7.1, p<0.05; Figure 3F). These findings imply that spine plasticity in S1 is related to texture discrimination learning.

#### SPINE LIFETIME AND SPINE SIZE

The reduction in spine elimination rate during texture discrimination resulted in an increase in spine lifetime (from t_1/2_ = 12 days to 38 days) for pre-existing spines present in both baseline imaging sessions (Figure 4A). New spines present in the imaging session immediately before training were not afforded the same protection from elimination and instead were eliminated at approximately the same rate (5.64 ± 1.45% per day) as they were formed (5.55 ± 1.45%) (F_(1,9)_=0.002, p=0.964). This finding suggests a specific preservation of pre-existing spines.

**Figure 4.**
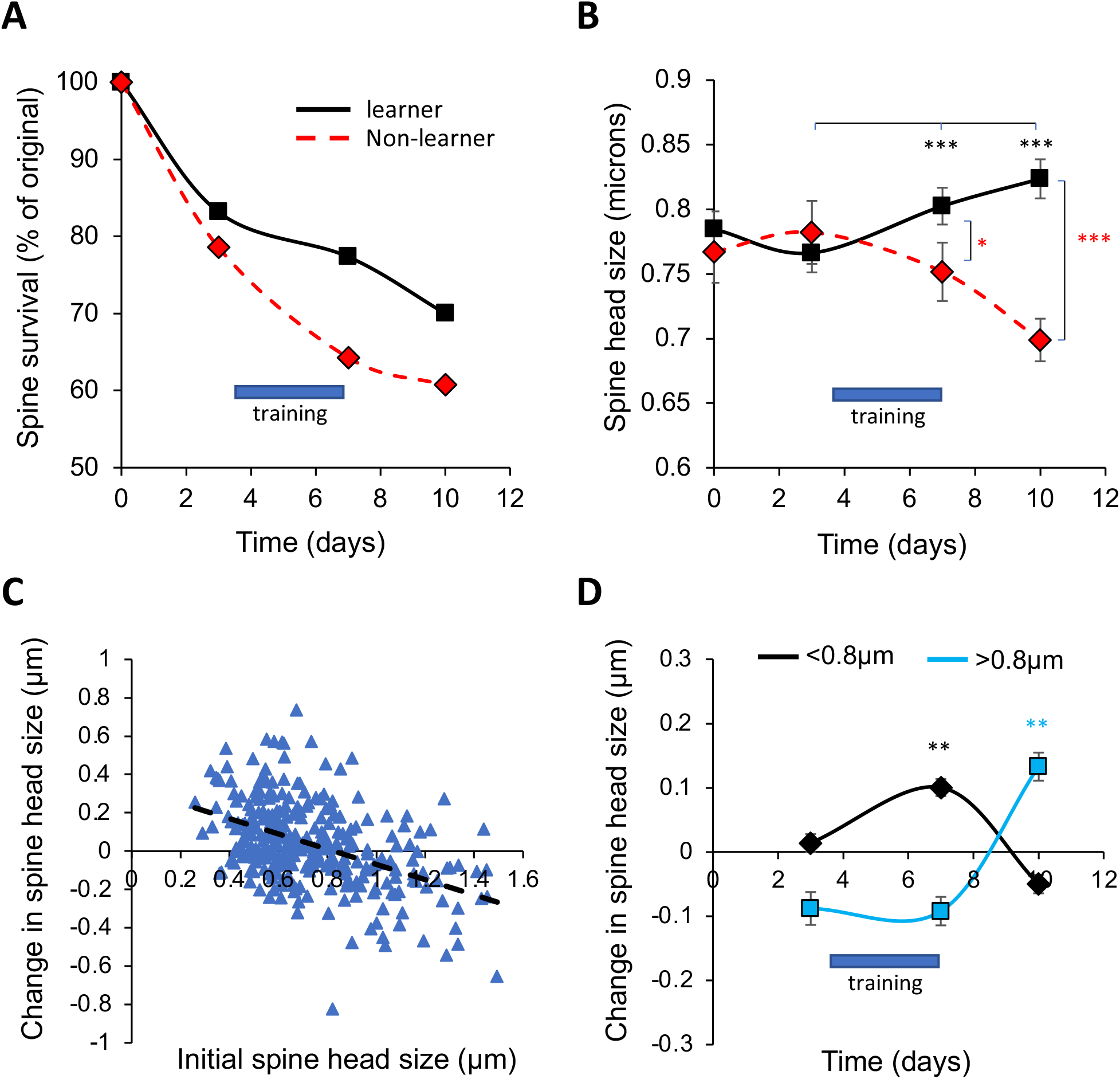
The effect of texture discrimination learning on basal dendrite spine lifetime and spine size. **A:** The number of spines present on a given day decays approximately exponentially in non-learners (red dashed line) but are preserved during training in learners (black line). **B:** Spine head size for the persistent spines (present on first and last imaging session) on basal dendrites. Spine size increases during training in learners (black) but in not non-learners (red). **C:** Changes in spine size are related to initial size. The point of least change occurs at approximately 0.8µm for the population of persistent spines. Regression line is shown (black dashed line). **D:** Difference in spine size between timepoints plotted at timepoints where the difference is measured (for example day 3 represents baseline 2-baseline 1). Spines greater than 0.8µm only increase post-training (blue). Spines smaller than 0.8µm only increase during training (black).

Previous studies have shown a close relationship between spine size and spine lifetime (Seaton et al., 2020; Yasumatsu et al., 2008). In concert with these observations, we found that the increased spine lifetime seen during texture discrimination was accompanied by a small but significant increase in spine head size for the stable population of basal dendritic spines that were present throughout imaging (Figure 4B), both immediately after texture discrimination (t_(354)_=2.86, p<0.002, matched pair t-test) and 3 days later (t_(354)_=3.38, p<0.0008, matched pair t-test). Due to spontaneous spine fluctuations, large spines tend to decrease and small spines tend to increase in size (Yasumatsu et al., 2008). For our data-set, the point of least change occurred close to 0.8µm (Figure 4C). Defining small spines as those with a head-width less than 0.8 µm, this population increased in size following texture discrimination (t_(200)_=7.53, p<0.0001) whereas during baseline it did not (t_(184)_=1.27, p=0.20). In the final imaging session, 3 days after the initial training period, the large spine population (>0.8 µm) increased in size (t_(118)_=2.75, p<0.003) (Figure 4D), despite the lack of training during the intervening 2 days.

For apical dendrites, no changes were seen in spine size across any of the six pairs of timepoints (e.g. baseline to post-texture discrimination t_(381)_=0.09, p<0.464 matched pair t-test). However, the animals that did not learn the discrimination showed approximately 9% smaller spine head size than learners in their baseline spine size measurements before training began (baseline 1: F_(1,404)_ = 5.00, p=0.0259, baseline 2: F_(1, 404)_= 4.83, p<0.0284). In this sense, the apical spine head size was predictive of learning outcome.

#### ODOUR DISCRIMINATION

We tested whether the effects on spine dynamics in S1 following texture discrimination were modality specific by measuring S1 spine dynamics in animals that performed an odour discrimination (see Methods). The relevant cue for reward was the odour of the sawdust in the bowl rather than the texture. This discrimination has many of the same task requirements as the texture discrimination, but the discrimination does not involve the somatosensory modality. Under these conditions, we did not find any changes in dendritic spine formation rates (F_(2,14)_=2.89, p=0.09), elimination rates (F_(2,14)_=1.8, p=0.21), nor on spine equilibrium (F_(2,14)_=2.36, p=0.14), either on apical or basal dendrites in S1 (2-way ANOVA for time-point and spine location F_(5,38)_ = 1.59, p = 0.19). However, spine head sizes in barrel cortex increased slightly on basal dendrites during training (t_(190)_=1.73, p=0.042), but unlike for the texture discrimination condition, this change did not persist to the probe-test time point three days later (Supplementary Figure 6D).

#### FORAGING WITHOUT A DISCRIMINANT

We also tested whether the effects seen during texture discrimination learning were related to the general increase in motor activity and exploration of the test arena. We imaged spines in animals that were allowed to forage in the testing arena without needing to discriminate between textures to gain a reward (Supplementary Figure 6). In this control condition, both bowls were of the same texture and both contained a reward and the animal’s access to the second bowl was not restricted after it had dug in the first bowl. In this condition, the mice performed all the same behaviours as those mice needing to discriminate between textures to obtain the reward, but were rewarded for digging irrespective of the nature of the outer surfaces of the bowls. Under these conditions, we did not find any changes in dendritic spine formation rates (F_(2,14)_=2.89, p=0.09), elimination rates (F_(2,14)_=1.8, p=0.21), spine equilibrium (F_(2,14)_=2.36, p=0.14) on apical or basal dendrites in S1 (2-Way ANOVA for time-point and spine location F(5,38)= 1.59, p-0.19). However, spine head sizes did change slightly between baseline and the end of the foraging period (Supplementary Figure 6H) and on average decreased in size (t_(73)_=2.08, p=0.0205), maintaining that decrease into the probe-trial 3 days later (t_(73)_=2.52, p=0.0069). This is the opposite direction to the change seen with texture discrimination learning, where spines increased in size.

We therefore conclude that texture discrimination is accompanied by changes in spine dynamics, spine lifetime and spine size in S1 cortex, and that these changes cannot be explained either by the accompanying increased foraging activity inherent in the paradigm, nor by the need to discriminate in general, but are specifically related to the need to discriminate a tactile texture to gain a reward.

### 5. Texture discrimination learning depends on the secondary somatosensory cortex

The secondary somatosensory cortex has been implicated in texture coding (Jadhav and Feldman, 2010; Jiang et al., 1997; Lieber and Bensmaia, 2019; Zuo et al., 2015). To test whether the secondary somatosensory cortex (S2) is required for learning a texture discrimination, we used the same strategy that we used for barrel cortex (S1) and injected floxed DREADD containing virus into S2 bilaterally 3-4 weeks prior to testing the mice on the texture discrimination assay, either with an activating i.p. injection of CNO or a control injection of saline. As the previous studies in S1 had indicated that CNO and GFP expression had no effect on learning, we did not repeat all combinations of control experiments in the S2 studies. However, we did compare CNO with saline injection in DREADD expressing animals as well as the effect of bilateral S2 inhibition on odour discrimination.

#### TEXTURE DISCRIMINATION

Examination of the DREADD expression locations showed that some injections produced expression entirely within S2 and some strayed into S1 (Figure 5A, B; Supplementary Figure 7). We therefore classified the cases where expression was exclusively located in S2 separately from those where expression overlapped with S1.

**Figure 5.**
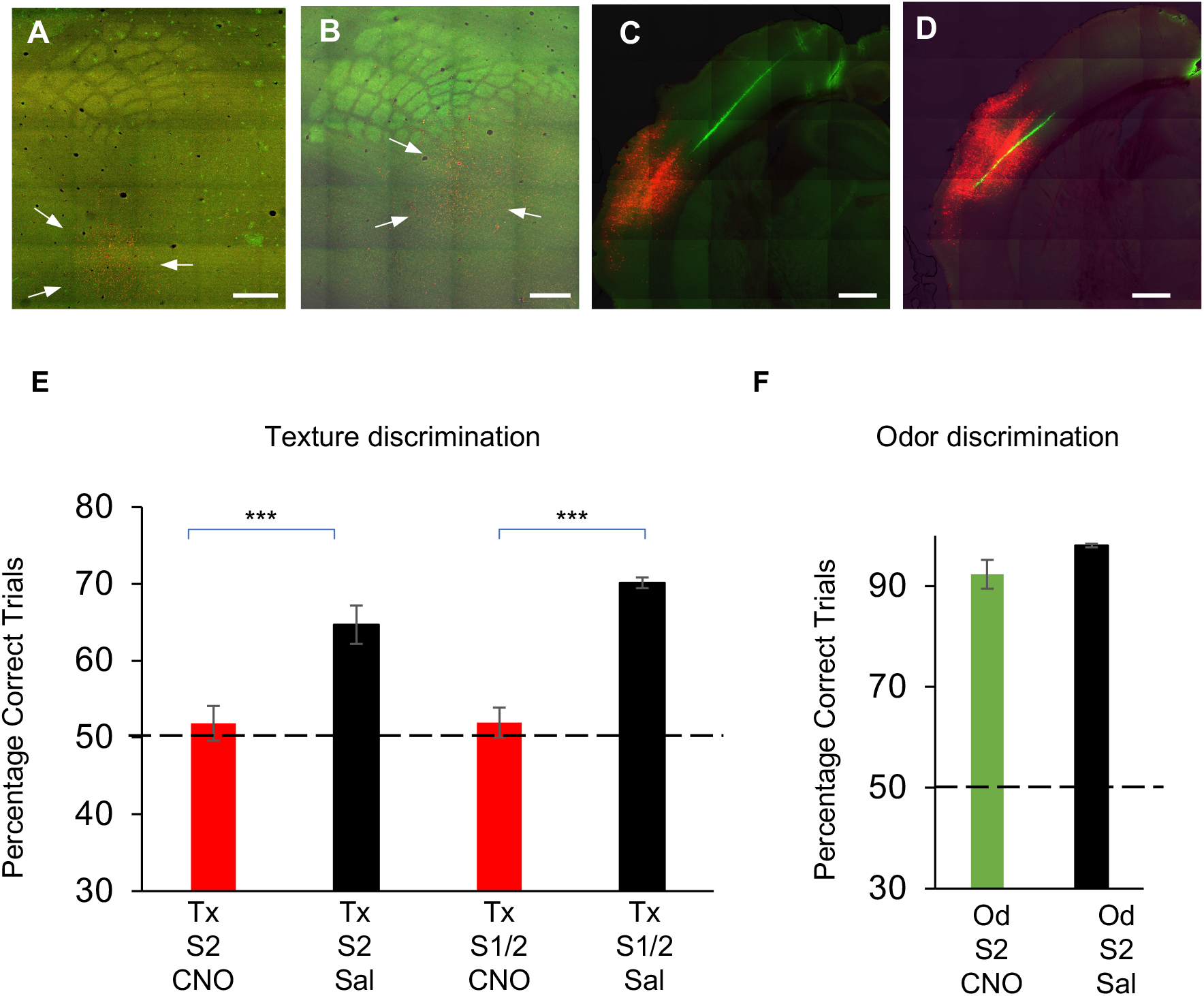
Effect of inhibition of S2 on texture and odour discrimination. **A:** Example of hM3D(Gq)-mCherry expression in S2 outside the barrel field. Arrows indicate expression (red). **B:** Example of expression of hM3D(Gq)-mCherry in S2 with some overlap into the barrel-field. **C:** Coronal section showing hM3D(Gq)-mCherry expression in S2 (red) and the Neuropixels electrode track (DiI coated). **D:** Adjacent coronal section to C showing electrode track (green) in S2 hM3D(Gq)-mCherry expression area (red). The identity of S2 in coronal sections was based on histology and electrophysiological responses to whisker stimulation. Scale bars 500 microns. **E:** Texture discrimination performance over 3 days was at chance levels when CNO was administered and hM3D(Gq) expression was only in S2 (Tx, S2, CNO), whereas animals learned when receiving saline instead (Tx,S2,Sal). In a few cases hM3D(Gq) expression was found in S1 and S2 (S1/S2) and the effect of CNO was again significantly different from saline. (*** p<0.005). **F:** hM3D(Gq) expression in S2 had no effect on odour discrimination when CNO was administered (green) and neither did saline injections (Od/S2/CNO or Sal). Scale bars 500µm.

We found that inhibition of S2 excitatory neurones had a significant effect on discrimination learning (Figure 5E). A two-way ANOVA involving DREADD location (S2 only versus S1 and S2) and treatment (CNO versus saline) was significant (F_(3,22)_=9.04, p<0.0006), and gave an effect of treatment on learning (F_(1,1)_ = 24,4, p<0.0001) but not location (F_(1,1)_ = 0.73, p=0.40). As can be seen in Figure 5, this was due to the hM3D(Gq)-CNO groups not learning the texture discrimination whether the injection was exclusively in S2 or some of the expression was also in S1. From our previous result (Figure 1) we would not expect learning to occur if some expression was in S1, though it is not clear that the small overlap with S1 of these primarily S2 injections would have been sufficient to prevent learning. However, in the cases where expression was exclusively located in S2 the mice did not learn either, showing that S2 is required for texture discrimination learning.

Post-hoc t-tests on just the S2 expression cases showed that there was a significant difference between saline controls and CNO treated animals (t_(14)_=3.58, p<0.0015). Scores for the mice in the hM3D(Gq)-CNO group tested for texture discrimination (Figure 5), did not differ significantly from chance levels of 50% correct (t_(2)_ = 0.74, p < 0.479; Cohen’s d = 0.247) while those in the saline group did (t_(2)_ = 5.88, p < 0.001; Cohen’s d = 2.22). In the saline treated cases 6/7 mice were judged to have learned the discrimination within 3 days by the binomial test (α=0.05), whereas in the CNO treatment group 1/9 mice learned the discrimination (Fisher’s exact probability = 0.0087, p<0.01). We therefore conclude that S2 is required for texture discrimination.

#### ODOUR DISCRIMINATION

As described previously for the S1 experiments, we wanted to test the possibility that inhibiting S2 activity had a non-specific effect that prevented learning or performance generally. Therefore, a different cohort of mice expressing DREADD in S2 performed the odour discrimination version of the foraging behaviour (see Methods). Of the 9 mice receiving bilateral injections of DREADD virus, expression was confined to S2 in 6 cases, while 3 also showed some S1 expression (all in the saline injected cases) and were excluded. We found that all of the mice learned the odour discrimination assay well. Scores for the mice in the hM3D(Gq)-CNO group (Figure 5F), differed significantly from chance levels of 50% correct (t_(2)_ = 13.17, p < 0.0057; Cohen’s d = 7.6). Mice in the hM3D(Gq)-Saline group also learned the odour discrimination (Figure 5F) and their scores differed significantly from 50% (t_(2)_ = 60.62, p < 0003; Cohen’s d = 34.99), and did not differ from the CNO treated group (F_(1,5)_=1.81, p=0.248). We therefore conclude that S2 cortex is not required for learning an odour discrimination and that the manipulation we used does not have a non-specific effect.

#### EFFECT OF DREADD ON NEURONAL ACTIVITY IN S2

Recordings were made in S2 with Neuropixel probes to measure the effectiveness of hM3D(Gq)-CNO on sensory responses. DiI was used to track the position of the penetration relative to the expression of the DREADD (Figure 5C, D). We found that sensory responses to single whisker stimulation (usually the C2 whisker) were strongly decreased in S2 (for 72.3% of excitatory cells and 36.4% of inhibitory cells) and slightly decreased for a smaller percentage of S1 cells (24.5% of excitatory cells and 22.2% of inhibitory cells) (Supplementary Figure 5). Network activity was also altered with the vast majority of neurones firing in phase with the delta-wave activity inhibited (Supplementary Figure 3B). In conclusion, activation of excitatory DREADD in inhibitory cells in S2 inhibited network activity and reduced sensory responses in most S2 neurones, with smaller effects on S1.

### 6. Second somatosensory cortex gates LTP in barrel cortex

The above experiments establish that S1 and S2 are both required for texture discrimination learning in mice and that changes in S1 spines occur during learning on basal but not for spine located on apical dendrites of layer 2/3 neurones. Recent studies have shown that S2 provides a projection to S1 that terminates strongly in layer I (Minamisawa et al., 2018). The S2 feedback projection is therefore positioned to engage the apical dendrites of L2/3 neurones where synapses appear unaffected by texture discrimination. The basal dendrites of L2/3 cells mainly receive columnar input from the L4 neurones in their barrel column (Hooks et al., 2011) and synapses on the basal dendrites do undergo structural plasticity (see section 4). We therefore hypothesised that a locally inert S2 input arriving on the apical dendrites of L2/3 neurones might gate plasticity on the basal dendrites of the same neurones. Therefore, we looked at the interaction between S1 columnar and S2 feedback connections converging onto layer 2/3 neurones located in S1 barrel cortex. The columnar input was activated electrically from the radially displaced L4 barrel and the S2 input was locally activated optically, using ChR2 expressed in the axons of S2 neurones (Figure 6). We used an LTP protocol that was sub-threshold for producing LTP from the columnar input alone. The stimulus protocol did not produce action potentials and so differed from spike timing dependent LTP (Hardingham et al., 2008). We compared the effect of in-phase stimulation of L4 and S2 with out-of-phase stimulation (see Methods).

**Figure 6.**
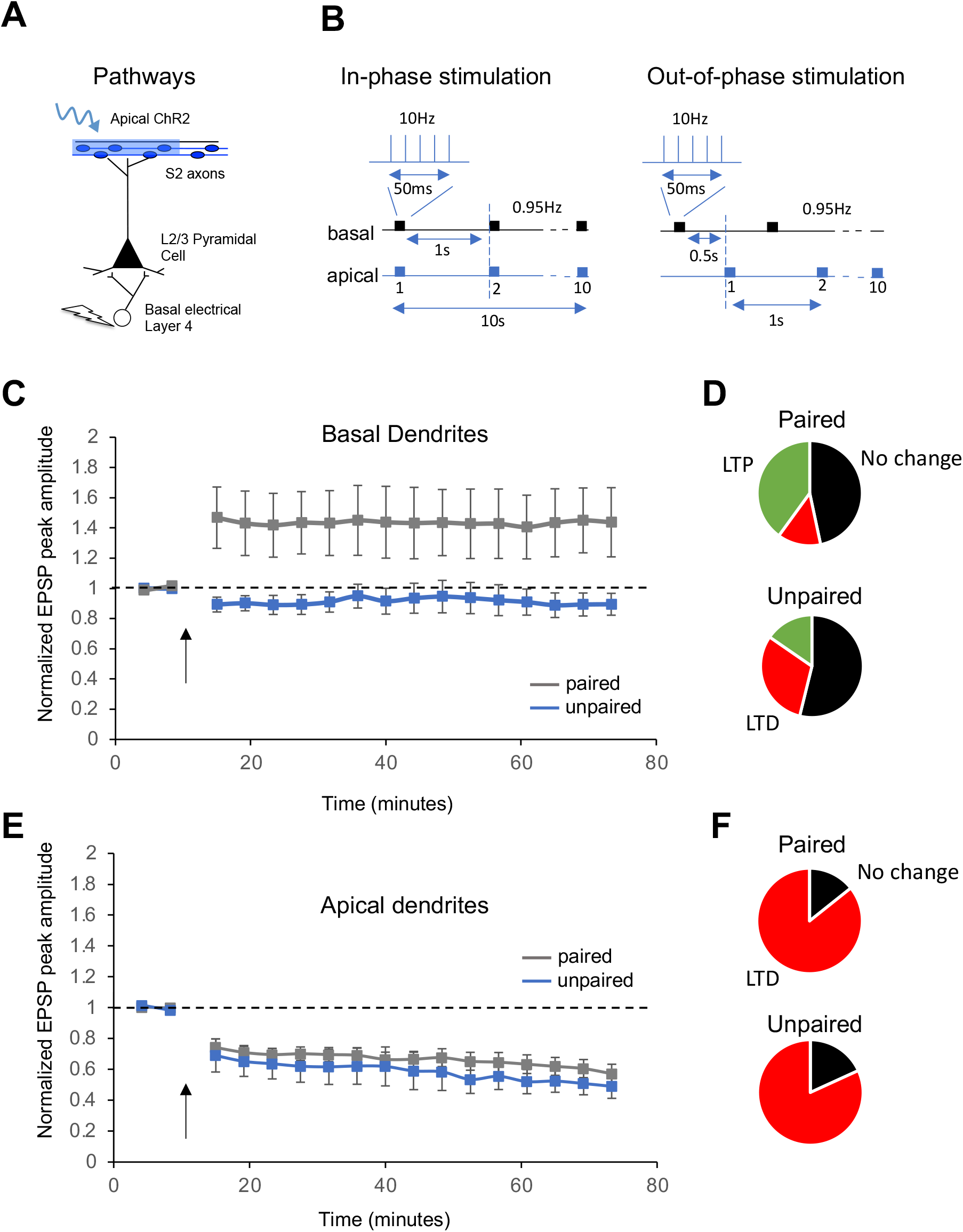
Apical gating of LTP in S1 by S2. **A:** Stimulus configuration for “Apical gating LTP pathways stimulated to induce LTP in layer 2/3 neurones. **B:** Left: In-phase stimulation timing diagram, showing bursts of 5 stimuli at 10Hz delivered one second apart and in-phase with S2 afferent stimulation optically. Right: As for the in-phase case, but apical and basal stimuli are separated by 0.5 seconds and therefore out-of-phase. **C:** Effect of in- and out-of-phase stimulus pairing on EPSP amplitudes in columnar Layer 4→layer 2/3 pathway. Normalised EPSP averages and standard errors in 4 minute epochs for basal stimulus response. **D:** Proportion of LTP (green) LTD (red) and cases where no change occurred (black) as a result of in-phase (paired) and out-of-phase (unpaired) protocols. **E:** Effect of in and out-of-phase stimulus pairing on gain of the S2 to S1 layer 2/3 pathway. Averages and standard errors in 5 minute epochs for basal stimulus response. **F:** Proportion of LTD (red) and cases where no-change occurred (black) as a result of in-phase (paired) and out-of-phase (unpaired) protocols.

#### L4 TO L2/3 PATHWAY (BASAL DENDRITES)

The general effect of in-phase L4 and S2 pairing was to produce an increased probability of LTP and a decreased probability of LTD at the layer 4 → layer 2/3 neurone pathway. The probability of LTP induction increased from 15 to 40% with apical gating. In several cases, in-phase pairing produced large changes in the synaptic gain of the basal inputs (maximum 387%). Conversely, out-of-phase stimuli produced a mildly depressing effect, causing LTD in just over 30% of neurones and having no effect on synaptic weight at all for the majority of cases (53%).

A two-way ANOVA looking at the effects of TIME (baseline vs 60 minutes post-pairing) and PHASE (in-phase pairing vs out-of-phase) was significant (F_(3,55)_=4.037, p=0.0118) with a significant effect of PHASE (F_(1,1)_=4.936, p<0.0307) and an interaction between PHASE and TIME (F_(1,1)_=4.862, p<0.0319). Post hoc t-tests showed that this was because the in-phase pairing protocol, where the apical and basal dendrites were stimulated simultaneously, differed from all other cases [in-phase differed from out-of-phase pairing (t_(26)_=2.21, p=0.0179), its own in-phase baseline (t_(28)_=1.99, p=0.0278), and the out-of-phase baseline (t_(26)_=1.86, p=0.0369)].

A similar conclusion is supported when each cell is compared with its own baseline. While the out-of-phase pairing does not lead to an increase in the response at 60 minutes post-pairing (t_(12)_=1.419, p=0.909), in-phase pairing causes a significant overall increase of 44 ± 22% (t_(14)_=-1.996, p=0.0329). The reason for the large variation around the mean for the in-phase pairing is that some responses increased by up to 387%, while others did not increase at all (6/15). The proportion of layer 2/3 cells that did not undergo either LTP or LTD was similar for in-phase and out-of-phase pairing protocols and also similar to values found in previous studies of the vertical columnar pathway from layer 4 to layer 2/3 (Hardingham et al., 2008; Hardingham et al., 2011). For the proportion of cells that are capable of undergoing LTP in the barrel cortex, in-phase pairing increases the probability of LTP at the expense of LTD (Figure 6).

#### FEEDBACK S2 TO S1 (APICAL DENDRITES)

The general effect of in-phase L4 and S2 pairing was to produce LTD at the S2 input. Out-of-phase stimuli also produced a depressing effect on the S2 to S1 layer 2/3 neurone pathway, with just over 80% undergoing LTD.

A two-way ANOVA looking at the effect of TIME (baseline vs 60 minutes post-pairing) and the effect of PHASE (in-phase pairing vs out-of-phase) was significant (F_(3,49)_=31.76, p<0.0001) with a highly significant effect of TIME (F_(1,1)_=94.31, p<0.0001), but not of PHASE (F_(1,1)_=0.98, p=0.34). Post hoc t-tests showed that this was because both in-phase and out-of-phase protocols produced a response at 60 minutes post-pairing that differed from baseline (in-phase t_(26)_=7.00, p<0.0001), and out-of-phase conditions (t_(20)_=6.66, p<0.0001)].

A similar conclusion is supported when each cell is compared with its own baseline. Both the out-of-phase pairing and in the in-phase pairing led to a decrease in the response at 60 minutes post-pairing (in-phase t_(13)_=6.97, p<0.0001; out-of-phase t_(10)_=6.70, p<0.0001). In-phase pairing causes a significant overall decrease to 58 ± 5.9% of control values (t_(13)_=6.97, p<0.0001) and out-of phase pairing causes a decrease to 49.9 ± 7.4% (t_(10)_=6.70, p<0.0001). The proportion of layer 2/3 cells that underwent LTD was similar for in-phase and out-of-phase pairing protocols (81% for out-of-phase and 85% for in-phase) (Figure 6).

#### Interaction of S2 and columnar S1 input

It is apparent that the plasticity of the columnar feedforward input onto layer 2/3 neurones located in S1 is affected by the timing of S2 stimulation. The probability of LTP increases and the probability of LTD decreases for the L4 input when input to the apical and basal dendrites is paired in-phase. In contrast, the S2 input to S1 simultaneously undergoes LTD in the same cells. The depression at the S2 input is not dependent on the potentiation of the columnar layer 4 input, because it occurs both for in-phase and out-of-phase pairing. This suggests that depression is not a homeostatic response to potentiation or vice versa. On the contrary, for cells that were stimulated out-of-phase, we found that the degree of depression is moderately positively correlated at the two sets of inputs (R^2^=0.50, F_(1,10)_=9.06, p=0.0147), that is to say, the more the cell was depressed at the S2 input the more it was depressed at the L4 input (Supplementary Figure 8). The correlation between changes in L4 and S2 inputs breaks down for in-phase stimulation; in this case, the two sets of inputs tend not to share the same direction of plasticity and neither was the degree of their opposite movement correlated (R^2^=0.063, p=0.388, F_(1,10)_=0.803). In fact, the L4 input overcame the naturally depressing effect of the stimulus protocol (low frequency of stimulus repetition without post-synaptic action potentials) and the tendency for the vertical columnar input within the cortex to to depress rather than potentiate (Hardingham et al., 2011). The potentiation that is “gated” by the S2 input is therefore both highly statistically and physiologically significant as it overcomes three tendencies toward depression.

## Discussion

### Primary and Secondary Somatosensory Cortex

We find that both S1 and S2 are required for texture discrimination learning in mice as bilateral inhibition of either cortical area prevents learning. It may not be entirely surprising that S1 is required for such learning as the discrimination is performed by the whiskers (Pacchiarini et al., 2020) and the barrel cortex is the major recipient of the somatosensory projection from the whiskers via the brainstem and the thalamus (VPm). Indeed, these findings are in agreement with previous studies in rat showing that barrel cortex ablation prevents whisker-based texture discrimination (Guic-Robles et al., 1992) and while detection of an object *per se* does not appear to require S1, more complex discriminations do (Ryan et al., 2022). However, S2 also receives a somatosensory thalamic input via POm, which theoretically could have bypassed the VPm!S1 route to enable learning. Indeed, we found that inhibition of S1 did not entirely abolish tactile responses in S2 to whisker stimulation (Supplementary Figure 5). Nevertheless, the residual S2 sensory activity was insufficient for learning when inhibition was increased in S1.

The finding that S2 was additionally necessary for texture discrimination (even though S1 remained active) is consistent with a number of its properties. First, it is known that S2 neurones respond to tactile textures (Jadhav and Feldman, 2010; Jiang et al., 1997; Lieber and Bensmaia, 2019; Zuo et al., 2015) and in humans, S2 even responds to images of textures in the absence of a tactile stimulus (Sun et al., 2016). Second, it is known that neuronal activity in S2 depends not only on the physical features of the tactile sensation, but also on salient aspects of the behavioural paradigm. For example, S2 neurones have been found to represent the difference between two independently presented tactile features when the reward depends on that difference (Romo et al., 2002) and to hold a trace of the first feature of the discriminated pair during encounter with the second (Salinas et al., 2000). In concert with this finding, S2 has been shown to be necessary for discriminating between different sequences of tactile stimuli (Bale et al., 2021). Finally, there is evidence to suggest that S2 neurones recall information in anticipation of touch (Condylis et al., 2020). The latter property of working memory and recall is highly likely to be important in the behavioural discrimination we have studied, where mice learn to reduce the errors they make by digging in the unbaited bowl by adopting a strategy of making several visits to each bowl before choosing (see Supplementary Material [Mouse texture discrimination assay.avi]).

### Structural Plasticity and LTP in S1

A major finding in the present study was that learning produced structural plasticity in S1. While several studies have shown that sensory deprivation can also cause structural plasticity in S1 (Seaton et al., 2020; Trachtenberg et al., 2002; Wilbrecht et al., 2010), there are rather few studies showing that learning has an effect (Kuhlman et al., 2014).

In contrast to the effect of sensory deprivation, we found that learning induced plasticity did not appreciably affect new spine formation, but rather induced plasticity in pre-existing spines, which increased in spine head size, producing a reduction of spine elimination and an increase in spine lifetime (Yasumatsu et al., 2008). Therefore, pre-existing connections were strengthened during learning rather than new connections being formed at a greater rate than normal. Spine size is correlated with AMPA receptor number and LTP increases AMPA receptor number in synapses (Kopec et al., 2007; Matsuzaki et al., 2004; Nusser et al., 1998; Shi et al., 1999). The results are therefore consistent with an LTP like increase in spine size. Spine size increases in barrel cortex are known to be αCaMKII-autophosphorylation dependent in S1 (Seaton et al., 2020) as is LTP itself (Giese et al., 1998), adding further credence to the idea that an LTP like process is involved. This finding is also consistent with studies showing that consolidation of fear conditioned learning is known to depend on LTP in the neocortex (Goto et al., 2021). In that study, cortical LTP was required on the second night’s sleep after induction of fear conditioning. In the present study, the enlargement of spines was present immediately after the last period of learning (where two periods of sleep would have elapsed), but then continued for two days (and therefore two periods of sleep) after that without any further training (Figure 4B).

For a minority of animals that failed to learn the discrimination, there was a decrease in pre-existing spine size rather than the increase seen with the animals that learned (Figure 4B). There was also a general correlation between S1 plasticity and performance at texture discrimination (Figure 3F), suggesting that the S1 plasticity was not trivial or incidental to learning. Neither did the plasticity occur due to increased exploration or generally environmental enrichment (Briones et al., 2004; West and Greenough, 1972), as animals that explored the same arena but were not required to discriminate in order to receive a reward did not show an increase in spine head size, but instead showed a decrease.

### Dendrite Specific Plasticity

It is likely to be important that the learning-induced plasticity was only found on basal dendrites of layer 2/3 neurones rather than on the apical dendrites. Apical dendrites have tended to dominate reports on plasticity in the neocortex, partly due to the ease with which they can be imaged. However, the apical dendrites predominantly receive feedback connections from other cortical areas including MI, S2 and peri-rhinal cortex (Schuman et al., 2021) rather than feedforward sensory information, which in the barrel cortex projects to the basal dendrites (Hooks et al., 2011). The lack of plasticity on the apical dendrites of mice that learned the discrimination suggests that feedback connectivity may play a modulatory role in learning rather than being modulated itself by the learning. This was borne out by our finding that S2 input can effectively gate LTP on basal dendrites of layer 2/3 neurones. When synchronously active, S1-L4 and S2 inputs conspired to produce LTP only in the basal dendrites not the apical dendrites. The degree of LTP produced by simultaneous apical and basal dendritic activation was far larger than that previously reported for LTP in the barrel cortex (Hardingham et al., 2003) and in one case reached 387% of baseline, suggesting a qualitatively different form of plasticity induction to that reported for neocortex before (Hardingham et al., 2003). Indeed, this type of neocortical LTP could also be produced in the absence of action potentials, which if applied in the living brain could enable neurones that were initially sub-threshold for responding to touch to be recruited to the active pool. This idea that remains to be tested, but if it were the case then it would increase the number of neurones responding to the stimulus and hence potentially aid discrimination.

Recent studies have shown that while S1 neurones primarily code for touch, they can also carry information that is usually associated with higher order cortical areas. For example, some cells can code for categorical information while others are lodged within the circuit for decision making (Buetfering et al., 2022). Furthermore, some cells, under the control of prefrontal cortex, are able to change their responsiveness dependent on the salience of the texture being discriminated (Banerjee et al., 2020). It is therefore possible that the plasticity we describe here for S1 is not simply involved in enhancing sensory coding, important though that is, but it may be involved in inducting plasticity in these smaller subsets of functionally important neurones in layer 2/3 of S1.

While our focus has been on the changes in basal dendrites, we did find one significant correlation between the size of the pre-existing population of spines on the apical dendrites (in the baseline condition before learning commenced) and future learning performance. Smaller apical dendritic spines were predictive of a lack of learning or at least a slower rate of learning. It is notable that a previous study on the effect of somatosensory learning on apical dendritic synapses (Kuhlman et al., 2014) showed that pre-existing spines on apical dendrites exhibit an increase in spine lifetime with learning that preceded and predicted an improvement in performance (Kuhlman et al., 2014). Since spine lifetime is generally known to be related to spine size (Figures 3 and 4) (Seaton et al., 2020; Yasumatsu et al., 2008), these findings suggest that apical dendritic spines may need to increase in size before learning can take place, possibly so that the synaptic gain in the feedback pathway is sufficient to gate plasticity.

### DREADD time course and memory consolidation

CNO is not predicted to have off-target effects at the dose used here (Jendryka et al., 2019), and we found that CNO injections in the absence of DREADDs did not prevent learning (Figure 1). The time course of CNO mediated DREADD action was fast enough to be active during the texture discrimination assay; on average it took 15 minutes to reach a maximum effect and the behavioural testing did not start until 30 minutes had elapsed. In recordings from anaesthetised mice, we did not see recovery of activity from a single i.p. injection of CNO (at a dose of 3.5mg/kg) within 2 hours (cases were not followed long enough for full recovery to be seen). Given that a single i.p. dose of CNO only produces detectable levels of CNO in the brain for less than 60 minutes, the continuing effect beyond 60 minutes was presumably due to the conversion of CNO to Clozapine (Jendryka et al., 2019; Manvich et al., 2018). In a Neuropixels recordings from a freely-moving rat, the CNO mediated DREADD inhibition effect could be seen to last for approximately 6 hours. Even allowing for faster metabolism of CNO and clozapine in mice, it is highly likely that the effect of the DREADDs lasted beyond the texture discrimination assay by up to 4 hours. Inhibition would therefore have lasted into the time-period immediately after learning and would have inhibited a proportion of the period when the animal would naturally have slept; both periods have been identified as important for the early stages of memory consolidation (Goto et al., 2021).

The effect of DREADD activation was not simply a complete inhibition of excitatory neuronal activity either in S1 or S2, though the effect was sufficient to prevent learning. While sensory responses were decreased during DREADD activation and abolished completely in some neurones, the largest effect of increased inhibition was to profoundly reduce network activity. The delta-wave activity characteristic of slow-wave-sleep was practically abolished and the burst-pause firing of neurones that is coordinated with the slow waves was strongly inhibited. The burst-pause firing of cortical neurones during anaesthesia and slow-wave sleep is known to be NMDA-dependent (Armstrong-James and Fox, 1988; Fox and Armstrong-James, 1986) and is highly sensitive to inhibition due to the voltage sensitivity of the NMDA channel (Mayer et al., 1984; Nowak et al., 1984). Given that sleep is important for learning and memory consolidation (Goto et al., 2021; Miyamoto et al., 2016), we cannot rule out the possibility that the DREADDs acted by altering cortical network activity necessary for consolidation during sleep rather than, or in addition to, its effect during learning.

Mice sleep more during the light-phase of the light-dark cycle, but sleep both during the light and dark-phases. Slow wave sleep episodes have durations that are bi-modally distributed with peaks at 5 seconds and 80 seconds (Soltani et al., 2019) and while episodes occur more in the light-phase (60-80%) than in the dark-phase (40%), they occur in both phases. Mice were housed in an environment with a 12 hour light/dark cycle and experiments were performed within 6 hours of the light-phase commencing. This means that the mice were returned to their home cages following texture discrimination training during the period when they were most likely to sleep with the DREADDs still active. Nevertheless, mice would have had a considerable period of the light-phase remaining for sleep after the effects of the DREADDs had worn off. Therefore, the effects we see, while not restricted to the learning period did not prevent activity during the entirety of the sleep period.

### Learning in Freely Moving Animals

One advantage of the present study is that the behaviour is entirely natural, in the sense that the mice are freely moving and able to forage in a test arena. As a consequence of exploiting this natural foraging behaviour of the animal, learning took place relatively quickly, with animals showing some learning on the first day (24 trials) and certainly by the third day (72 trials). This is in contrast to head-fixed paradigms where the animal usually requires hundreds of trials per day over 1-2 weeks to learn a detection or discrimination (Gilad and Helmchen, 2020; Kuhlman et al., 2014; O’Connor et al., 2010). With head fixed trials, quite a large component of the learning is related to the animal understanding the timing and sequence of the discrimination, which often involves auditory cues to start or wait and time-windows within which the reward is available. With freely moving behaviour, the animal makes its own decisions about when to perform different actions, and in that sense has less to learn, particularly given that for mice foraging behaviour is innate. The plasticity that takes place in freely moving animals may therefore also be more closely related to that which takes place naturally in the somatosensory cortex.

Freely moving animals are also likely to be less stressed than head-fixed animals. In this study the mice underwent a two-week period of habituation before the experiment began to minimise stress levels. Corticosteroid levels are initially high during the early stages of head fixation (Juczewski et al., 2020) and are known to inhibit plasticity (Daw et al., 1991). The current methodology may therefore be useful where rapid learning is required.

### Network Topology and Learning

Finally, neural network circuits have made use of back-propagation to modify the synaptic weights of the input layer in the circuit based on the output of the circuit (le Cun, 1988). There has been interest in this mechanism for implementing predictive coding, where the feedback connections carry information predictive of what might be received at the input layer and the error between the two sources modifies synaptic connection weights to improve the prediction (Bastos et al., 2012). The difficulty has often been to justify the applicability of such models to the way learning occurs in the brain because a physical manifestation of back-propagation process is unlikely (Crick, 1989) and currently remains undiscovered. In the present study, we have found evidence not of back-propagation of an error signal, but potentiation in a lower cortical area dependent on gating from a higher order cortical area. However, the plasticity described here offers a plausible mechanism by which feedback information from higher order cortical areas might influence feedforward information without employing back-propagation. Our studies have shown that apical dendrites in S1 receive feedback information from S2, which then modifies feedforward information onto the basal dendrites of the same S1 neurones. One further line-topological feature of this type of network plasticity is that the information is coded in the projection to the input cortical area rather than (as could still be the case) in its projection to the higher order area. While we have not ruled out plasticity in the S1→S2 pathway, we have provided evidence that the S2→S1 pathway modifies the sensory input pathway to S1.

## Methods

### 1. EXPERIMENTAL MODEL AND SUBJECT DETAILS

#### MOUSE

All the procedures were carried out in accordance with the Animal (Scientific Procedures) Act 1986. A total of 171 mice were used in the study across genotypes and experiments. All the mice used were either WT C57Bl/6Jax (RRID: IMSR_JAX:000664, acquired from Charles River, UK) or transgenic homozygous PV-Cre strain, kindly gifted by Prof. Jack Mellor (University of Bristol, UK). The genotype of PV-Cre homozygous mice was determined using PCR analysis with DNA obtained from ear biopsies. All the dendritic spine imaging experiments and most of the behavioural training experiments were performed on C57Bl/6-Jax mice. A subset of behavioural training experiments was performed on homozygous PV-Cre mice. The mice were 9 to 13 weeks old when behaviour or imaging experiments were initiated, while synaptic plasticity experiments were performed on mice aged 10-14 weeks.

#### RAT

Additionally, recordings were made from one Lister Hooded rat aged 4 months, with a chronically implanted Neuropixels electrode (IMEC, Leuven, Belgium). Recordings took place over a period of 8 days and generated approximately 6 hours of data each day (results from 2 days of recordings are reported here).

### 2. Behavioural training

Mice were trained on a texture discrimination in which they were free to move around the testing apparatus as described before (see Pacchiarini et al, 2020). Food restricted mice were trained to dig for a food reward hidden in sawdust (Lignocel hygienic animal bedding, Lignocel, Germany) contained in circular plastic bowls. The 3-D printed bowls had either a smooth outer texture or a vertically grooved rough texture. The internal texture of the bowls was smooth and identical. One pellet of Chocorice (glucose syrup and cocoa powder (5.5%) coated rice flakes (76.5%), Crownfield, UK) was used as a food reward in each trial. Food restriction started 5 days before day 1 of behavioural training, which was aimed to gradually bring down the animal’s body weight to 87-90% of its original weight. The bowls were placed in two chambers separated by a compartmental partition (Figure 1, Supplementary Figure 1). Both chambers were accessible from a common holding area when the door was opened. Water was available ad libitum throughout the behaviour in the holding chamber. Animals were handled for two weeks prior to starting the training. The behavioural assay comprised 4 phases over 8 days.

**Phase 1 Habituation:** was carried out for 2 consecutive days during which the mice were exposed to the arena for 3 trials. In between trials, they were restricted to the holding compartment. In the first trial, mice were free to move throughout the empty arena for 10 minutes. During the next 2 trials, mice were free to explore the arena for 5 minutes, this time with rewarded bowls with the same outer texture present in both the chambers. The texture used during habituation became the rewarded texture in all subsequent phases of the assay.

**Phase 2 Training and test trials:** were run on two consecutive days after day 2 of habituation. There were two components; first, 4 consecutive trials were run where the mice were free to explore both the chambers. One chamber contained a rewarded bowl and other chamber an unrewarded bowl with a different texture. Each of these preliminary training trials lasted for at least 5 minutes, or in some instances, a given mouse was given extra time to make sure it dug in both the bowls. Second, 24 consecutive testing trials were run divided into 6 blocks of 4 trials each. Test trials were the same as the preliminary training trials except that once the mouse had made a choice to dig, it was isolated with its chosen bowl (with digging being interpreted as the measure of choice). During this period they could not reach the other chamber. Individual testing trials lasted for at least one minute or until the mouse had dug in at least one bowl. If they did not dig in either of the bowls after 5 minutes had elapsed, the trial was aborted and the result was recorded as an abstinence and not considered further in the analyses.

**Phase 3 Further test trials:** The two-day training and testing phase was followed by a further testing day in which the preliminary trials were omitted and there were 24 testing trials with just one baited bowl. All other details were the same as the previous days 2 days.

**Phase 4 Follow-up:** For the next two days, mice were left in the home cage still under food restriction. On the last day of the experiment, “Phase 3 Further test trials” was repeated. If an imaging session was performed on the same day as the behavioural training, it was performed after the behavioural experiment the same day.

**Randomisation, counterbalancing and other factors:** The position of the bowls was pseudo randomly changed from trial to trial such that the probability of having a rewarded bowl in any given location was 0.5. In each experiment, half of the subjects in a cohort were presented with the reward in a smooth and the other half in a grooved bowl. To make sure the experimenter was blind to the location of the mouse with respect to the chambers, the holding area was obscured until the guillotine doors were opened to release the mouse into the chambers. The experimenter was blind to the location of DREADD injection, as the injection locations were only confirmed from histology *post hoc*.

The experiments were performed in a room dimly lit with red light. To conceal any odour cue produced by the reward, the sawdust used in the bowls was mixed with fine powder of the Chocorice used as a reward (2% of the sawdust). In a subset of experiments, Clozapine N-oxide dihydrochloride (CNO) was injected before the behavioural experiment began. In these cases CNO (HelloBio, UK) was injected intraperitoneally (3.5mg/Kg of the subject’s bodyweight) 30 minutes before the start of experiment. Control experiments were identical except that Saline was injected instead of CNO. CNO was injected only on day 3 to day 5 of the behaviour experiments (i.e., phase 2 and phase 3 described above). To ascertain the effect of CNO on behavioural performance some of the mice expressing GFP in area S1 were trained on the texture discrimination with an equivalent CNO injection, while some other mice without any viral infection in the brain were also trained following similar CNO injection as described in the results section.

**Odour discrimination:** The odour discrimination experiments were performed similarly, except that instead of mixing 2% chocorice powder with the sawdust we mixed in an odourant. Both the bowls had the same external texture, but one bowl was filled with sawdust mixed with 0.5% ginger powder (Stonemill, UK), while other bowl was filled with 0.5% cinnamon powder (Stonemill, UK); again, only one of the bowls was rewarded. Other details of the procedure were the same as for the texture discrimination.

### 3. Surgery

All surgeries were performed on mice aged 6 to 10 weeks. For dendritic spine imaging a mixture of flexed GFP and CamKII-Cre AAV virus was injected in the barrel cortex, while for DREADD mediated S1 or S2 inhibition, AAV-DREADD virus was injected in the respective areas. In the subset of experiments designed to study the effect of S2 silencing during the texture discrimination learning on S1 structural plasticity, AAV-DREADD virus was injected in S2 while ipsilateral S1 was injected with GFP-Cre virus mix, combined with an optical window implantation over the barrel cortex. All the injection coordinates were determined based on a standard mouse brain atlas (Paxinos and Franklin, 2008). The following procedures were common in every surgery: briefly, before surgery, mice were given an intramuscular injection of Colvasone (0.4 mg/kg, Norbrook, UK) and Metacam (0.5mg/kg, Boehringer Ingelheim, Germany) for pain and inflammation management and a 1ml intraperitoneal injection of Saline (0.9%) and Glucose (4%) to manage dehydration during surgery. Lidocaine (100 µl, 2%, Hameln pharmaceuticals, UK) was injected subcutaneously at the incision site for local anaesthesia. Deeply anaesthetised mice (1.5-2.0% isoflurane in 1.5 L/minute medical oxygen) were head-fixed in a stereotaxis frame (model 963, Kopf Instruments). Body temperature was maintained with a thermostatically controlled heating pad. After shaving the fur, the head was disinfected with Videne (Ecolab, UK) and 70% Ethanol. All viral injections were performed using beveled glass micropipettes with a tip diameter of 10-20 µm fitted to an Ultra-micro syringe pump (WPI, USA), and a Micro4 controller (WPI, USA). Following surgery, the mice recovered in a warm recovery chamber in dim light until ambulatory and were then transferred to the home cage and housed in normal 12hr dark-light cycle with food and water provided ad libitum until the food restriction started (where applicable). Post-surgery the mice were single housed without any additional enrichment in the cages other than handling tubes and nesting material.

#### TRANSCRANIAL WINDOW IMPLANTATION AND INTRACRANIAL rAAV VIRUS INJECTIONS

To achieve sparse labelling of neurones with GFP, a mixture of flexed AAV GFP (pAAV-FLEX-GFP-Virus, titer ≥ 1×10¹³ vg/mL, Addgene 28304-AAV PHPeB) and CamKII-Cre (pENN.AAV.CamKII 0.4.Cre.SV40, titer≥1×10¹³ vg/ml Addgene 105558-AAV9) virus was injected in S1 barrel cortex. 250nl of the virus solution (CamKII Cre-AAV 1×10^9^ vg/mL, in equal proportion with Flex-GFP 1×10¹^2^ vg/mL and mixed with 10% Fast Green for visualization) was injected (25nl/min) at 2 locations into the layer 2/3 of barrel cortex. After making an incision in the scalp, the periosteum was cleared, and the outer skin layers were adhered to the dry skull with tissue adhesive (Vetbond, 3M). S1 injection locations were marked as indicated in Table M1. A surgical stainless-steel head-plate was implanted with dental cement (Prestige Dental, Super Bond C_B kit) keeping the injection locations approximately at the center of the window. Mice were then head-fixed using the steel head-plate just implanted. A 3-mm-diameter craniotomy was performed using a micro-drill. The skull was removed gently. The dura was also removed carefully. Throughout the surgery the brain was covered with cortex buffer. The injection micropipette was lowered with a micro-manipulator (Kopf Instruments) to 250 µm below the surface. The glass micropipette was left for a further 5 minutes in the brain after injection had finished. A sterile 3 mm glass coverslip concentrically glued to a 5mm glass coverslip was placed over the exposed area and sealed with Super Glue and dental cement as described earlier (Goldey et al., 2014) Imaging began at least 3-week post recovery period as described previously (Crowe and Ellis-Davies, 2014)

#### DREADD INJECTION FOR S1 OR S2 SILENCING

For experiments to test the effect of S1 or S2 silencing on texture learning the DREADD virus was injected in S1 or S2 layer 4. For DREADD mediated silencing of S1 or S2, excitatory DREADDs were expressed in inhibitory neurones. For this purpose, pAAV-hDlx-GqDREADD-dTomato-Fishell-4 (Addgene, 83897-AAV9) viruses were injected in C57Bl6-Jax mice or in some cases in rats. The ‘DREADD-Fishell’ viruses are expressed in most types of inhibitory neurone and once activated by CNO increase their activity and thereby decrease activity in the excitatory neurones (Figure 2, Supplementary Figures 2, 3, 4, 5). For these injections a small incision was made over the area of interest and small holes were drilled at the injection locations using microdrill. ‘DREADD-Fishell’ virus was injected in C56-Bl6 Jax mice at the locations and dilutions given in the Table M2 for S2 injections. In the case of S1 injection, DREADD-Fishell virus was injected at the locations and at the dilutions given in Table M3. In some cases, DREADD-flexed (pAAV-hSyn-DIO-hM3D(Gq)-mCherry, Addgene 44361-AAV9,) was injected in PV-Cre mice at the same dilutions and locations as described above for C57Bl/6 Jax mice. Since there was no difference in behavioural or electrophysiological outcome of the CNO injection between the PV-Cre and C57Bl/ 6-Jax mice the data were pooled and analysed together.

#### CHANNELRHODOPSIN INJECTION IN S2 FOR SYNAPTIC PLASTICITY EXPERIMENTS

For Synaptic plasticity experiments C57Bl/6-Jax mice were injected with AAV-Chronos (pAAV-Syn-Chronos-GFP, Addgene 59170-AAV5) virus in area S2 at locations and concentrations given in Table M2.

### 4. 2-photon Imaging of dendritic spines

#### DENDRITIC SPINE IMAGING

Dendritic spines were imaged at 4 time points across the duration of the experiment (Figure M1). For each ROI, two baseline time points (B1 & B2) were imaged two days apart, and third imaging time point (T1) three days after the second baseline, and a follow up imaging time point (F1) was two days after T1 without behavioural testing in-between. T1 and F1 imaging sessions were both carried out after behavioural testing. Spines were imaged as described earlier (Seaton et al., 2020). During imaging, the animals were lightly anaesthetised with Isoflurane (1.5-2.0%) mixed with medical oxygen. The mice were head fixed with implanted stainless-steel head-plate. After each imaging session the animals were injected with 800µl of saline and returned to the home cage after a brief recovery period. The imaging was performed on an Olympus BX68 microscope using 25x water-immersion objective lens (Olympus, W Plan-APOCHROMAT,1.05 numerical aperture), 6 mm galvo mirrors, and a beam expander to fill the back aperture. PrarieView software was used for data acquisition. The Ti:sapphire two-photon laser (Chameleon Vision S, Coherent, USA) used for imaging was mode locked at 900 nm with power at the back aperture in the range of 10-30mW. The emission wavelengths were band-passed between 525–570 nm with an IR filter included in the light path. The section of dendritic branch was imaged as z-stacks with interframe interval of 0.5 µm and digitally magnified to the pixel size of 0.091 µm with 1024x1024 pixels in a field of view. The apical dendrites were imaged from layer 1 and upper layer 2/3 between 50 to 150 µm depth below the dura. Only secondary or tertiary branches were imaged. For the basal dendritic spine imaging the dendritic branches were generally selected by tracing them to their somatic origin points, in some cases where the origin point of the branches were not clear, the branches were considered as basal dendrites if some cell bodies were found at the same depth and the dendritic branch was mostly restricted to a similar depth.

**Figure M1:**
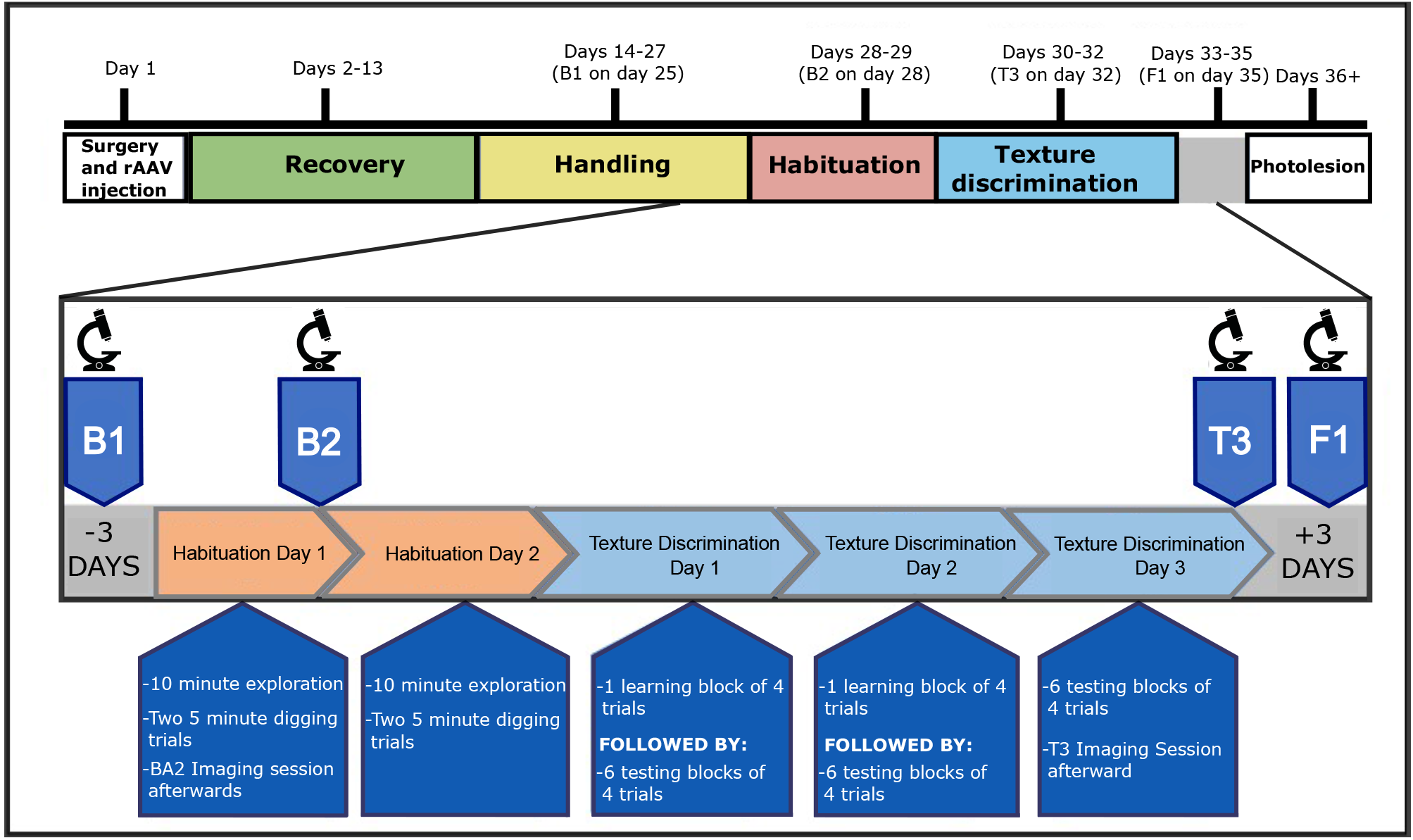
Timeline of experiments. A schematic illustration of the experimental time-line for spine imaging and texture learning in mice. Handling of mice started approximately 2 weeks after surgery. One week after the start of handling, they were put on food restriction and slowly in the next one week their body weight was brought down to 87%-90% of pre food restriction body weight. The first imaging sessions, labelled here as B1 (Baseline 1) and B2 (Baseline 2), formed the baseline period, while T1 (Testing 1) and F1 (Follow up 1) formed the 2 imaging time points after training. The zoomed in version of habituation and texture learning is showing details of behavioural training program. In some cases, mice were trained only for 5 blocks of 4 trials. Refer to the ‘behavioural training’ section above for details.

#### ANALYSIS OF DENDRITIC SPINE IMAGES

Image J was used to analyse all the z-stack images following procedures described earlier (Holtmaat et al., 2009; Seaton et al., 2020). Raw images were deconvolved using Fiji Deconvolution lab plugin (Sage et al., 2017) using point spread function generated for the same imaging configuration. Only protrusions that were at least 0.4µm from the dendritic shaft were considered as dendritic spines (Holtmaat et al., 2009). Relatively flat sections of dendrites with signal to noise ratio greater than 4 were marked in all corresponding images taken on different days, and individual spines were annotated and tracked across the imaging sessions. Spine formation and elimination rates were calculated by dividing the number of gained or lost spines respectively at each time point by the number of spines on the dendrite on the first imaging session. For the formation or eliminations rates, the numbers of spines were divided by number of days elapsed between the two imaging sessions. By scrolling across the z-axis of the z stacks, the spine head width, neck width and neck lengths were measured for each spine at the frame of best focus. The spine head width was taken as the greatest diameter across spine head. The spine head sizes follow a log normal distribution when measured this way and it is similar as describe earlier with a different method (Loewenstein et al., 2011). In some analyses only “Always Present” (persistent) spines were considered, while in others all spines were considered, as described in the Results. The analyses were performed by several different analysts blind to the behavioural training outcome, the injection locations and/or the hypothesis under consideration.

#### PHOTOLESION

Photolesions were made in the imaging area and identified post-mortem from cytochrome oxidase stained sections through the barrel field (Wong-Riley, 1979). In brief, the mice were deeply anaesthetised with isoflurane and head fixed under the 40X objective lens (Olympus, WPlan-APOCHROMAT 0.8NA, water). The laser was mode locked to 800nm and power was adjusted to 50-65mW. After locating where spine imaging was performed, laser was focused at 400 µm from brain surface to lesion layer 4. Galvos were centered and the shutter was opened for 10-12 minutes. Mice were then perfused with PBS followed by 4% paraformaldehyde under terminal anaesthesia using urethane.

#### CYTOCHROME OXIDASE LABELLING OF BARRELS

The perfused cortex was flattened by carefully pressing the fixed brain between two slides (having first removed the thalamus and striatum) and stored overnight in 4% paraformaldehyde containing 20% sucrose. The flattened brain was then transferred to PBS containing 20% sucrose which was later tangentially sectioned at 35 µm followed by cytochrome oxide staining as described earlier (Wong-Riley., 1979). In a subset of animals these sections were stained for vGLUT2 instead of Cytochrome oxidase for in order to identify the barrels. For locating the injections of DREADDs the images of these stained sections were overlaid on a standard flattened cortical map (Gamanut et al., 2018). The Red fluorescent DREADD injection locations were marked on this map with respect to barrels.

### 5. In vivo Electrophysiology

*In vivo* electrophysiological measurements were performed to quantify the pharmacokinetics of DREADD activation using CNO. Acute in vivo electrophysiology was performed in mice to quantify optimal concentration of CNO, and minimum time required to silence excitatory activity via DREADD activation of inhibitory neurones. Chronic *in vivo* electrophysiological recordings of spontaneous activity were performed in awake freely moving rats for 10 hours using neuropixel probes (IMEC, Belgium) to quantify the duration of S1 after a single injection of CNO in animals expressing DREADDs in interneurones. The details of both these methods are given below.

#### ACUTE IN VIVO ELECTROPHYSIOLOGY IN MICE

Anaesthesia was induced with isoflurane (Piramal Critical Care UK, 4% in O_2_) and maintained with an i.p injection of urethane (1.5g/10 mL, Sigma-Aldrich, USA) mixed with 0.1 mL of Acepromazine Maleate (2 mg/mL, Elanco, UK) at a dose of 1.5 mg/g body weight, at an anaesthetic depth equivalent to Guadel stage III-2. An initial dose of 70% was administered and topped up as judged from indicators of anaesthetic depth. Anaesthetic state was monitored by spontaneous cortical activity, hind limb withdrawal reflex and respiratory rates during the experiment (Friedberg et al. 1999). Body temperature was maintained at 37°C by a heating pad controlled by a rectal thermistor. Anaesthetised mice were secured in a Narishige SR-6 stereotaxic frame (Narishige International, London, UK); the skull was thinned over a 2 x 2mm area above the barrel-field centerd on the D1 barrel (at approximately 3mm lateral to the midline and 1.5 mm caudal to bregma). A small fleck of thinned skull was removed just large enough to permit entry of the microelectrode.

For single unit recordings in S1 barrel field, we used carbon fibre micro-electrodes (Carbostar-1, Kation Scientific, Mn USA) as described before (Fox et al., 1980). Penetrations were aimed at an area that was close to the virus injection areas, usually between two virus injection sites where the blood vessels permitted. The electrode recording was amplified and filtered (600-6KHz) before spikes were discriminated based on their amplitude between an upper and lower amplitude threshold (Neurolog, Digitimer, UK).

For multi-site recordings in S1 and S2, we used Neuropixels 1.0 probes (IMEC, Belgium). An oval area of the skull was thinned (major axis mediolateral (ML) 2mm, minor axis anterior-posterior (AP) 1mm) location for probe implantation centered on (in mm): ML: 1.75, AP: 1.5, +/- 0.2. A small area of thinned skull was reflected to allow entry of the probe at an angle of either 20 or in other experiments 38 degrees away from the midline. The AP coordinates of the penetration were determined by aiming the electrode at the location where we could see bur holes from the virus injection surgery. The electrode’s narrow edge opposed the cortical surface.The Neuropixel probe connected to the computer (running OpenEphys (v 0.5)) via the acquisition chassis of the National Instruments board (National Instruments, USA) was inserted at approximately 2 µm per second to the target depth of 3.5 mm.

Following the insertion of the probe, a piezo-electric whisker stimulator was used to deflect the whisker (Physik Instrumente, UK)(Fox et al., 2018). The working principal whisker was chosen based on the peak response to stimulation assessed by the LFP, unit sensory response and the number of channels with putative sensory response. The validity of the principal whisker was assessed from histology post-mortem using the lesion location when using carbon fibre micro electrodes, but it was not possible with neuropixels probes, which recorded across many principal whiskers. Stimuli were applied at either 1Hz or 0.2Hz.

For carbon fibre micro electrode recordings, individual neurones were isolated by making small adjustments to the recording depth. Only a single cell was recorded per animal due to the effective irreversibility of the CNO effect when administered i.p., (at least within the time available for recording in an acute preparation). A period of 30 minutes baseline was acquired before injecting the CNO and a period of at least one hour was followed beyond the injection time.

For neuropixels recordings we waited one hour for adjustment period following probe insertion. A waiting period was not necessary with the carbon fibre electrodes. For neuropixels, three consecutive 60 minute sessions were recorded: 1) baseline without stimulation, 2) principal whisker stimulation, 3) principal whisker stimulation following an i.p injection of Clozapine N-oxide (CNO) hydrochloride at a dose of 3.5mg/kg (100mM, HelloBio, UK).

#### CHRONIC IN VIVO ELECTROPHYSIOLOGY IN RATS

Rats were implanted with a Neuropixel probe during a stereotaxic surgery (David Kopf Instruments, USA), under Isoflurane anaesthesia (0.5-4% Piramal Critical Care, UK). In addition, all animals received antibiotics (intraperitoneal injection of Baytril, 0.85 mg/kg, Bayer, Germany) and analgesia (subcutaneous injections of Metacam, 1 mg/kg, Boehringer Ingelheim, and Buprenorphene, 0.05 mg/kg, Ceva, France), immediately after anaesthesia induction. The probe was implanted into the barrel cortex (AP: -1.9 mm, ML: -4.2 mm, at an angle of 15 degrees away from the midline), at a rate of 2 µm/s, until it had penetrated 3.50 mm into the brain. Two screws positioned above the cerebellum served as an animal ground, which were connected to the probe ground and reference. Following implantation, the probe shank outside of the brain was sealed with paraffin wax and then covered with bone cement (Zimmer Biomet, UK). The probe ground and reference were also connected to a copper mesh wall surrounding the implant, which was shaped into a protective housing and further covered with cement.

Electrophysiological recordings in freely moving rats: The recordings were performed within a small open field arena and a sleep box. The rat (n=1) was first trained to forage for food pellets in an open field and habituated to a sleep box over 3 days, prior to the start of the experiment. Following a 7-day recovery period, the rat was food restricted and then underwent a series of recording sessions during exploration and rest. On each experimental day, they were placed in the open field arena where the animal performed pellet chasing for 15 minutes. Afterwards, the animal was transferred to the sleep box until >15 minutes of quiet rest data was collected. Then, the rat received an intraperitoneal injection of either saline or CNO (3.5 mg/kg, HelloBio, UK) and immediately returned to the sleep box for 30 min. This was followed by an additional open field (15 min) and sleep box recording. The animal was then placed in the start box every hour, between 2 and 7 hours after the time of injection. Following this, (at ∼7 hours 15 minutes) the rat returned to the open field for the last recording.

Analysis of firing rates during exploration/rest: We compared changes in population firing rates after injection with Saline (day 1) and CNO (day 2). Firing rates were measured separately for each cell (n-=47) in 5-minute time segments, in each behavioural session. The first 15 minutes were selected from all behavioural sessions, except the rest period immediately after DREADD, in which the first 30 minutes were selected. This enabled us to align the data recorded in the saline and CNO recording days. The time series of neuronal firing rates on each day were then smoothed (Gaussian kernel, 1 SD=1 time bin). The baseline rate on the CNO and saline days was calculated by taking the mean rate of the three segments recorded during the first open field session of the day, for each cell. Thereafter, the firing rate in each 5-minute segment was expressed as a normalised score for each cell (rate - baseline / rate + baseline), and averaged across the population.

#### ELECTROPHYSIOLOGICAL RECORDINGS: DATA ANALYSIS

Individual recordings were concatenated, such that saline and CNO data were processed together for the freely moving animal data (see below), while periods of whisker stimulation and network activity could be analysed together for the anaesthetised mouse data. Spike sorting on the concatenated data was performed using Kilosort 2.5 (Pachitariu et al., 2016), after which the resultant spike clusters were manually curated using Phy 2 (Rossant and Harris, 2013). Spike clusters were assigned to the channel with the largest voltage trough-to-peak amplitude (VTP), measured on the cluster average spike waveform. Only units with a clear refractory period (+-0.15 ms) were kept for further analysis. The resultant time series were then analysed through Matlab and C software, produced in-house.

To distinguish spike clusters into excitatory and inhibitory cells, we separated neurones based on the shape of the spike waveform and its tendency to burst. First, spike width was measured by calculating the time between trough and peak (TtoP) from the average spike waveform of each cluster (Bartho et al., 2004; Pala and Stanley, 2022; Sofroniew et al., 2015). Second, we defined “burstiness” (or tendency to fire bursts of spikes) using a method described from Kim et al. (2012) (Kim et al., 2012). Briefly, each neuron’s spike autocorrelation [+-20ms, 1ms bins] was calculated and burstiness expressed as the ratio between counts in 1-6ms bins, divided by counts across the whole histogram. Clusters with TtoP ≤ 0.5ms and burstiness ratio ≤ 0.4 were defined as inhibitory cells and the remainder as excitatory neurones.

Response to sensory stimulation under Anaesthesia: Each mouse underwent continuous 0.2Hz whisker stimulation under urethane anaesthesia, before and after the CNO injection, as described above. Responses to stimulation were estimated by generating a peri-stimulus time histogram (PSTH). PSTHs were constructed with 1ms bins in a 50ms window, after stimulation. Sensory responses were defined as the firing rate in the 3 - 50 ms post-stimulus bins. Those cells with a rate lower than 0.5 Hz during this window both before and after CNO treatment were excluded from further analysis. To normalize changes in sensory response across cells in the population, we calculated a sum-difference ratio before and after DREADD activation (firing rate after CNO – firing rate before CNO/ firing rate after CNO + firing rate before CNO).

**Change in spontaneous firing rates after DREADD activation:** Spontaneous firing rates were calculated in three-second time segments between stimulations. The first and last second of the inter-stimulus interval was removed from the analysis to prevent the influence of whisker deflection on spontaneous rates. The resultant firing rates were then used to calculate a change in rate score [(Mean rate after CNO - Mean rate before CNO)/Sum of activity before and after CNO)].

**Spectral analysis:** To investigate the effect of DREADD on delta power (0.5-4 Hz), we selected the LFP channel in S1 and S2 that showed the largest response to whisker stimulation. Delta power was then calculated in consecutive 10-second windows during the recording (both before and after DREADD activation) using the Welch function (Hamming window; 50% overlap).

### 6. In vitro electrophysiology

Recordings were made from 14 C57Bl6-Jax mice aged 10-14 weeks. Mice were killed by decapitation at 3-4 weeks after injection of rAAV Chronos virus in S2 as described earlier. The brain was quickly removed after decapitation and immediately placed in ice-cold slicing solution (in mM: 108 choline-Cl, 3 KCl, 26 NaHCO_3_, 1.25 NaH_2_PO_4_, 25 D-glucose, 3 Na Pyruvate, 1 CaCl_2_, 6 MgSO_4_, 285 mOsm, bubbled with 95% O_2_ 5% CO_2_). Coronal slices were cut at 350µm thickness in ice-cold slicing solution using a vibrating microtome (Microm HM650V). Slices were then transferred to a holding chamber containing normal ACSF (in mM: 119 NaCl, 3.5 KCl, 26 NaHCO_3_, 1 NaH_2_PO_4_, 2 CaCl_2_, 1 MgSO_4_, 10 D-glucose, 300 mOsm bubbled with 95% O_2_ 5% CO_2_). Slices were incubated at 37^0^C for 30 minutes, then returned to room temperature before recording. The barrel cortex was identified using a mouse brain atlas (Paxinos and Franklin, 2008) and detection of visible striations in layer 4 corresponding to barrels. Whole cell voltage/current clamp recordings were obtained from L2/3 neurones using borosilicate glass electrodes (4-7 MΩ) filled with a potassium-gluconate based internal solution (in mM: 110 K-gluconate, 10 KCl, 2 MgCl_2_, 2 Na2ATP, 0.03 Na2GTP, 10 HEPES, pH 7.3, 270 mOsm). Series resistances were compensated for and input resistance were measured during the recording to ensure recording stability.

Postsynaptic potentials were evoked by two means. The first was by electrical stimulation with a monopolar stimulating electrode (Tungsten, 0.5MΩ Harvard), placed in L4, in a direction radially below the recorded cell. Extracellular stimuli consisted of 1ms current pulses set to produce a 4-6mV monosynaptic EPSP in the postsynaptic neuron. For optical stimulation, a 473nm laser pulse was applied through the objective over the cell in a horizontal line parallel to the pia using a Rapp UGA-42 Firefly point scanning device (Rapp Optoelectronic, Germany) for localized photomanipulation (Kohl et al., 2011). The power of the optical stimulus was adjusted to produce a 4-6mV monosynaptic EPSPs recorded in the postsynaptic neuronal soma.

In the whole-cell current-clamp configuration, the LTP experiments consisted of a 10-minute baseline period during which electrical and optical EPSPs were evoked with a separation of 1 second and repeated at a frequency of 0.1Hz. This was followed by 5 minutes of paired stimulation where the electrical stimuli were paired with optical stimuli in bursts of 5 at a frequency of 20Hz in 5 groups of 10 bursts where electrical and optical stimuli were given at the same time. We then recorded for a further 60 minutes using the same stimuli timings as during baseline, so normalized LTP levels could be calculated for optical as well as electrical inputs (Figure 6). Electrophysiological data were acquired using a using Signal software (CED) low pass filtered at 1-3kHz (Digitimer, UK) and sampled at 20KHz for off-line analysis on a computer (RM, UK). Both electrically and optically evoked EPSPs were measured using an automated routine that compared a window in the baseline membrane shortly before the EPSP with the peak EPSP amplitude.

### 7. Cfos methodology

#### IMMUNOHISTOCHEMISTRY (IHC)

Details of the immunoreagents used are located in Table M4. Following completion of the behavioral assay, mice were left in a dark room in their home cages for 90 minutes to allow maximal expression of the cFos protein. They were then given a lethal dose of pentobarbital (Euthatal, Boehringer Ingelheim Animal Health UK Ltd.) and immediately perfused transcardially with 0.1M phosphate buffered saline (PBS; pH 7.4), followed by 4% paraformaldehyde (PFA) in PBS. The brains were removed and for a tangential view of cortical layers, the cortex was dissected and flattened between two glass slides (Lauer, Schneeweiß, Brecht, & Ray, 2018). The sample was then post-fixed for 24 hours at 4 °C in 4% PFA and equilibrated in PBS containing 25% sucrose at 4 °C. Fixed brains were then cut tangentially into 35 μm sections using a freezing-microtome (Leica Biosystems SM200 R) and sections stored at -20 degrees in a cryoprotectant solution (50% sucrose, 1% polyvinyl pyrrolidone and 30% ethylene glycol in 0.1M PBS). For the IHC, individual floating sections were thoroughly rinsed in PBS solution and blocked for 1 hour with 2% goat serum and permeabilized with 0.05% Triton X-100 in PBS (PBST). The slices were then incubated at 4 °C for two days with a mixture of the following antibodies: rabbit anti-cFos polyclonal primary antibody (1:5,000; Synaptic Systems), guinea-pig polyclonal anti-VGluT2 primary antibody (1:2,000 Synaptic Systems). To confirm the cell type infected with DREADD, a second subset of DREADD injected slices were also incubated in rabbit anti-PV polyclonal antibody (1:2,000; Swant Inc.) and incubated at 4 °C for 24 hrs. After incubation, slices were washed thoroughly in PBST and incubated in a solution of Alexa Fluor 647-conjugated anti-rabbit (1:1,000; Abcam) and either Alexa Fluor 488 conjugated anti-guinea pig antibody (1:500; Abcam) or Alexa Fluor 568-conjugated anti-guinea pig antibody (1:500; Abcam) in the 2% blocking solution for 2 hours at room temperature. The second batch of anti-PV slices were incubated in a solution of Alexa Fluor 488-conjugated anti-rabbit antibody. All slices were then washed in PBST and incubated in DAPI (1:15,000; Sigma Aldrich) in PBS for 10 minutes. Slides were washed in PBS, air dried and mounted in Fluoromount Aqueous Mounting Medium (Sigma Aldrich) and cover-slipped using Vectashield (Vector Laboratories). The following control solutions were used for both protocols: (1) a solution without the primary antibody, (2) a solution without the secondary antibody and (3) a solution without any antibody. Compared with the normal solution, no fluorescence was detected (data not shown).

#### IMAGING AND ANALYSIS OF COLOCALIZATION

Tissue was visualized using a confocal laser scanning microscopy (Zeiss LSM880) and the location of fluorescent neurones outlined using Adobe Photoshop (CS4) and analysed using Imaris (Bitplane) Image Software. Brain regions were outlined by comparing the slice images with the corresponding atlas maps (Paxinos & Watson, 1998; Wang, Sporns, & Burkhalter, 2012). Once the area was measured, Fos-immunopositive neurones were counted using the automated Dot Quantification Analysis feature of Imaris. As with previous studies, cFos neurones were counted only when clear immunostained nuclei were co-localized with DAPI staining (Oshitari, Yamamoto, & Roy, 2014; Yokoyama et al., 2013). DREADD infected neurones were readily visualized with native fluorescence from the mCherry expression in the PV-Cre mice. One representative section per brain region from each mouse was used for quantification, including both hemispheres.

In order to compare the cFos activity in specific regions within and outside the injection area, a new channel was created which only included colocalized cells (either DREADD/GFP and cFos signal present). Cells were detected using pixel intensity threshold (mCherry: 49337.14, GFP: 32639.50, cFos: 32639.50) whereby all pixels with a value higher than the threshold value are classified as feature pixels, and pixels with a lower value classified as background pixels. These thresholds were used to generate the new channel which only contained the colocalized voxels and excluded channels outside the region which exhibited no correlation (Costes et al., 2004). The colocalized channel was manually inspected to ensure background had not been included, if this occurred, the thresholds were altered to correct for it. Following the creation of the colocalization channel, the Imaris spot detection feature was used to detect spots within a region of interest (ROI) of either 400 x 400μm or 200 x 200μm. Within the ROI, the mean density (cells/mm2) was calculated as the number of colocalized cells in one region divided by the mean area size of that region (Lin et al., 2018). In addition to this, a ‘minimum area of super-threshold’ filter (4.15μm) and a ‘background subtraction’ filter was applied, this adds a Gaussian filtered channel filtered by ¾ of the spot radius, the intensity center of the spot is then used to detect the spot for the channel of interest (Costes et al., 2004). All spot detection images were manually inspected to ensure the background was not being registered, as a result, some spots were manually removed or the spot detection thresholds altered in sections which contained a large amount of background noise.

To confirm the chemogenetic activation of PV cells, two DREADD injected cases were stained against PV. Two ROIs of 1000 x 1000μm area were defined (one encompassing the DREADD injection sites and one with no DREADD injection viral spread). Imaris was used to automatically detect the colocalization in the two channels (DREADD and PV). As before, colocalization was determined using a pixel intensity threshold (mCherry: 979.46, PV: 664.17, DAPI: 45600.00). The same ‘quality’ and ‘background subtraction’ filter was applied to remove noise signal and the spot detection feature used to count cells.

### 8. Quantification and statistical analysis

Details of the statistical analysis are included where they appear in the results section, however some general principles are noted here. Statistical analysis was performed using JMP 16 software (SAS Software, USA). Distributions were tested for normality before applying parametric statistics. Spine head size data was found to be log-normally distributed, and log transformed before applying parametric statistical methods. If the data was not normally distributed non-parametric statistics were used. For parametric tests, ANOVAs were run to find effects and interactions before using post-hoc t-tests. For behavioural data, we used a binomial test to gauge whether individual animals learned, in addition to parametric tests to ascertain whether groups of animals learned under different conditions. Linear regression was used to test the strength and statistical significance of correlations. Statistical significance was indicated by p<0.05 and lack of significance for p>0.05 (actual p values are indicated at the relevant locations in the text, e.g. p>0.05, p<0.05, p<0.03, p<0.02, p<0.01, p<0.005, p<0.003, p<0.002, p<0.001, p<0.0002, p<0.0001). Matlab, R and Sigma Plot software were used for data analysis and plotting graphs.

---

**Table M1:** Location and viral load for S1 layer 2/3 injections for spine imaging.

| Stereotactic coordinates |  |  | Viral load |  |
| --- | --- | --- | --- | --- |
| Posterior from bregma (mm) | Lateral from midline (mm) | Ventral from dura ($\mu\text{m}$) | Volume (nl) | Dilution |
| 1.0 | 3.0 | 250 | 250 nl of viral mix | 1:1 mixture of AAV-GFP ($1 \times 10^{11}$ vg/mL) and AAV CaMKII-Cre ($1 \times 10^9$ vg/mL) |
| 1.1 | 3.0 | 250 | 250 nl of viral mix | 1:1 mixture of AAV-GFP ($1 \times 10^{11}$ vg/mL) and AAV CaMKII-Cre ($1 \times 10^9$ vg/mL) |

**Table M2:**
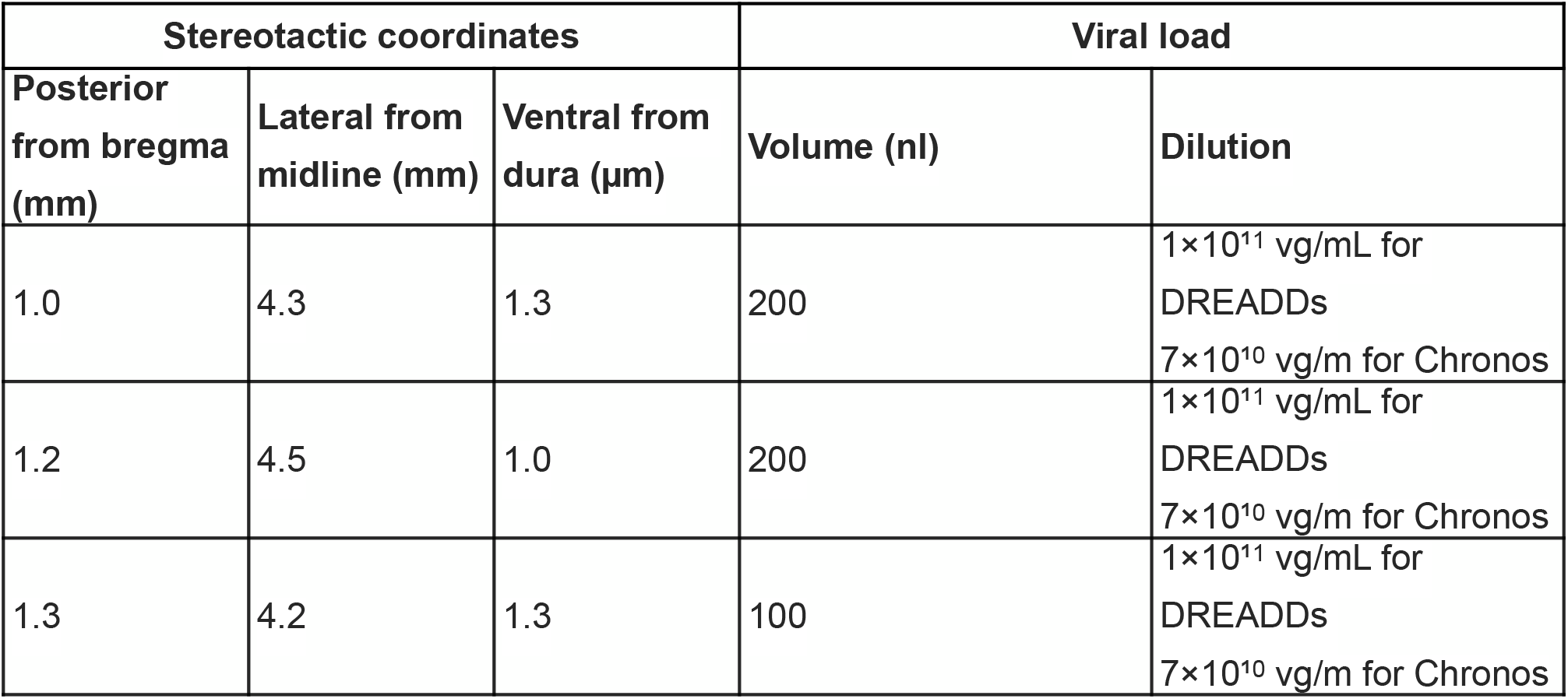
Location and viral load for S2 injections of DREADDs or Chronos for LTP experiments.

| Stereotactic coordinates |  |  | Viral load |  |
| --- | --- | --- | --- | --- |
| Posterior from bregma (mm) | Lateral from midline (mm) | Ventral from dura ($\mu\text{m}$) | Volume (nl) | Dilution |
| 1.0 | 4.3 | 1.3 | 200 | $1 \times 10^{11}$ vg/mL for DREADDs<br>$7 \times 10^{10}$ vg/mL for Chronos |
| 1.2 | 4.5 | 1.0 | 200 | $1 \times 10^{11}$ vg/mL for DREADDs<br>$7 \times 10^{10}$ vg/mL for Chronos |
| 1.3 | 4.2 | 1.3 | 100 | $1 \times 10^{11}$ vg/mL for DREADDs<br>$7 \times 10^{10}$ vg/mL for Chronos |

**Table M3:**
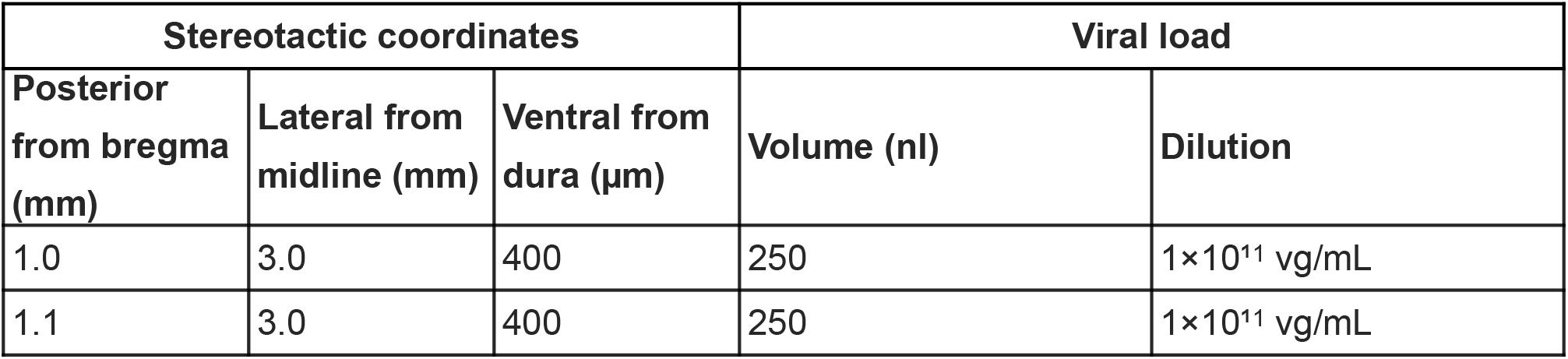
Location and viral load for S1 injections of DREADDs for behavioural testing

**Table M4:** Anitbodies used for cfos immunohistochemistry.

| **Antibody** | **Dilution** | **Source** | **Identifier** |
| --- | --- | --- | --- |
| Rabbit polyclonal anti-cFos | 1:5,000 | Synaptic Systems | 226 003 |
| Guinea-pig polyclonal anti-VGluT2 | 1:2,000 | Synaptic Systems | 135 404 |
| Rabbit anti-PV polyclonal | 1:2,000 | Swant Inc. | PV 27 |
| Alexa Fluor 647-conjugated anti-rabbit | 1:1,000 | Abcam | ab150079 |
| Alexa Fluor 488 conjugated anti-guinea pig antibody | 1:500 | Abcam | ab150185 |
| Alexa Fluor 568-conjugated anti-guinea pig antibody | 1:500 | Abcam | ab175714 |
| Alexa Fluor 488-conjugated anti-rabbit | 1:500 | Abcam | ab150077 |

