## Supplemental Figures 1-8 for "Interdependence of primary and secondary somatosensory cortices for plasticity and texture discrimination learning"

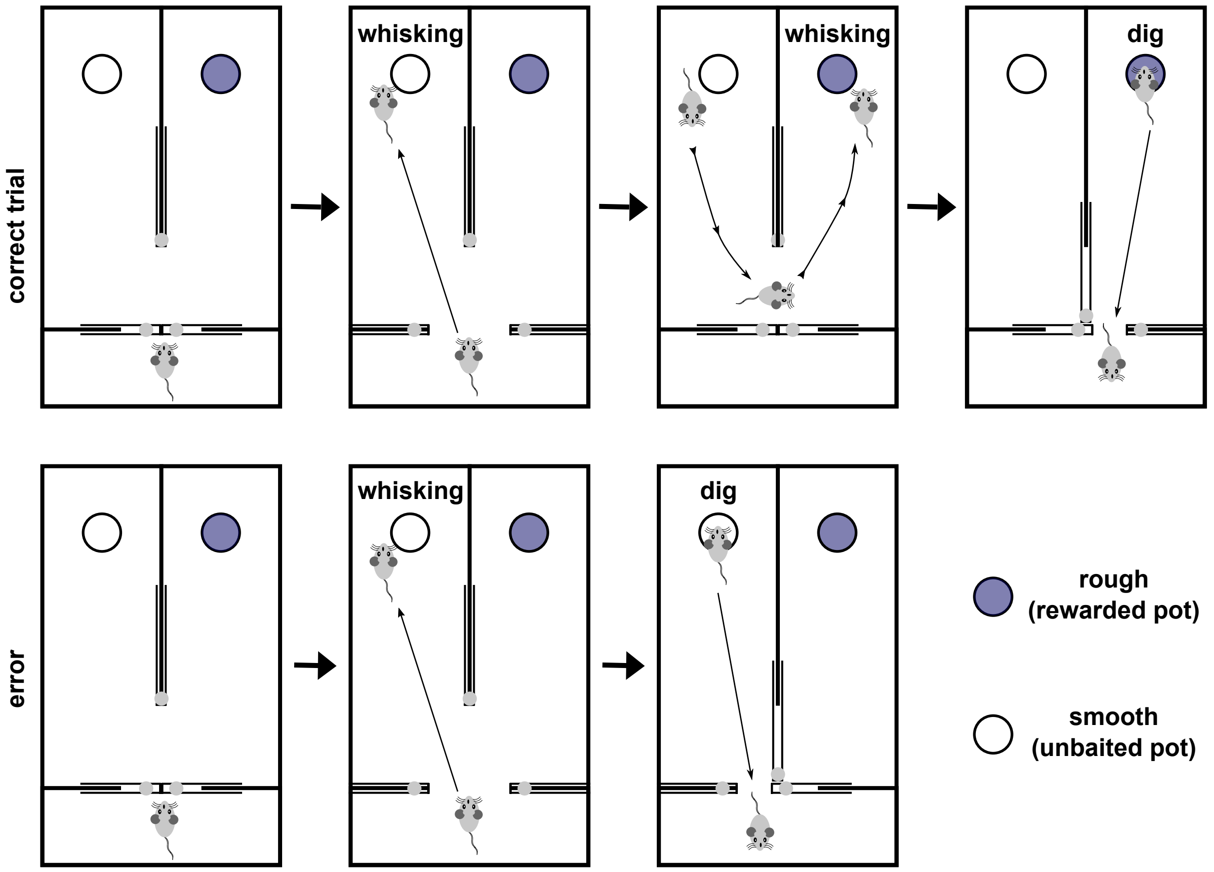


**Supplementary Figure 1**: The arena for texture discrimination and illustration of correct and incorrect choices. **Top:** a correct trial is illustrated. Once the mouse is allowed into the foraging space it can move freely between the two pots in the two compartments. In this case, the animal chooses to dig in the sawdust filled pot on the right, which is scored as a correct choice as it contains the food reward (cocoa pop, mauve circle). When the animal digs, the door to the neighbouring compartment is closed so that the mouse cannot dig in both pots. The door to the holding area is also opened at this point so that the animal can return. The holding area also contains a water bowl. The correct bowl is coded by the texture on the outer surface of the pot. Rough and smooth bowls are pseudo-randomly placed in left and right compartments in the trial sequence so that the reward is equally likely to be found in either location. **Bottom:** An incorrect trial is illustrated. When the animal digs in the unbaited bowl (white), access to the neighbouring compartment is closed and access to the holding area is opened.

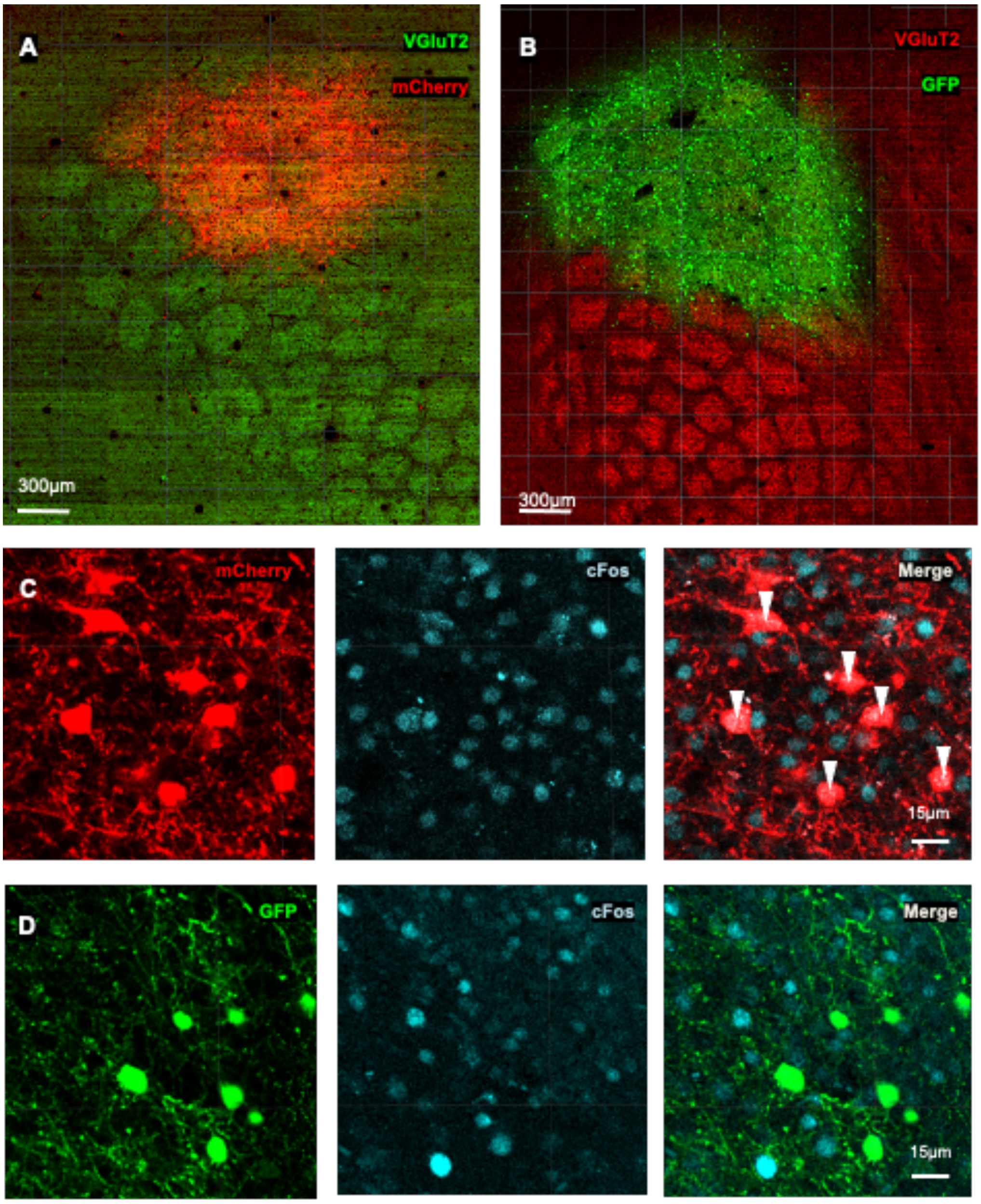


**Supplementary Figure 2**. Cfos staining in PV and non-PV cells in the barrel cortex. **A:** The vector containing the DREADD sequence also contains mCherry which can be seen here to be expressed in the caudal barrel field (red). The thalamic axons are visualised with VGlut2 antibody (green). **B:** In control experiments GFP was used instead of DREADD. GFP expression in PV cells (green) is shown against the barrel field thalamic axon staining using VGlut2 (red). **C:** left - PV cells expressing mCherry. Middle - cfos staining in the same area. Right - The merged image shows bright cfos staining in the PV cells and the contrasted non-PV cells cfos staining. **D:** Similar to C except with GFP staining. Left - GFP staining in PV positive neurones. Middle - cfos staining in same area. Right - merged image shows PV and non-PV cell cfos staining.

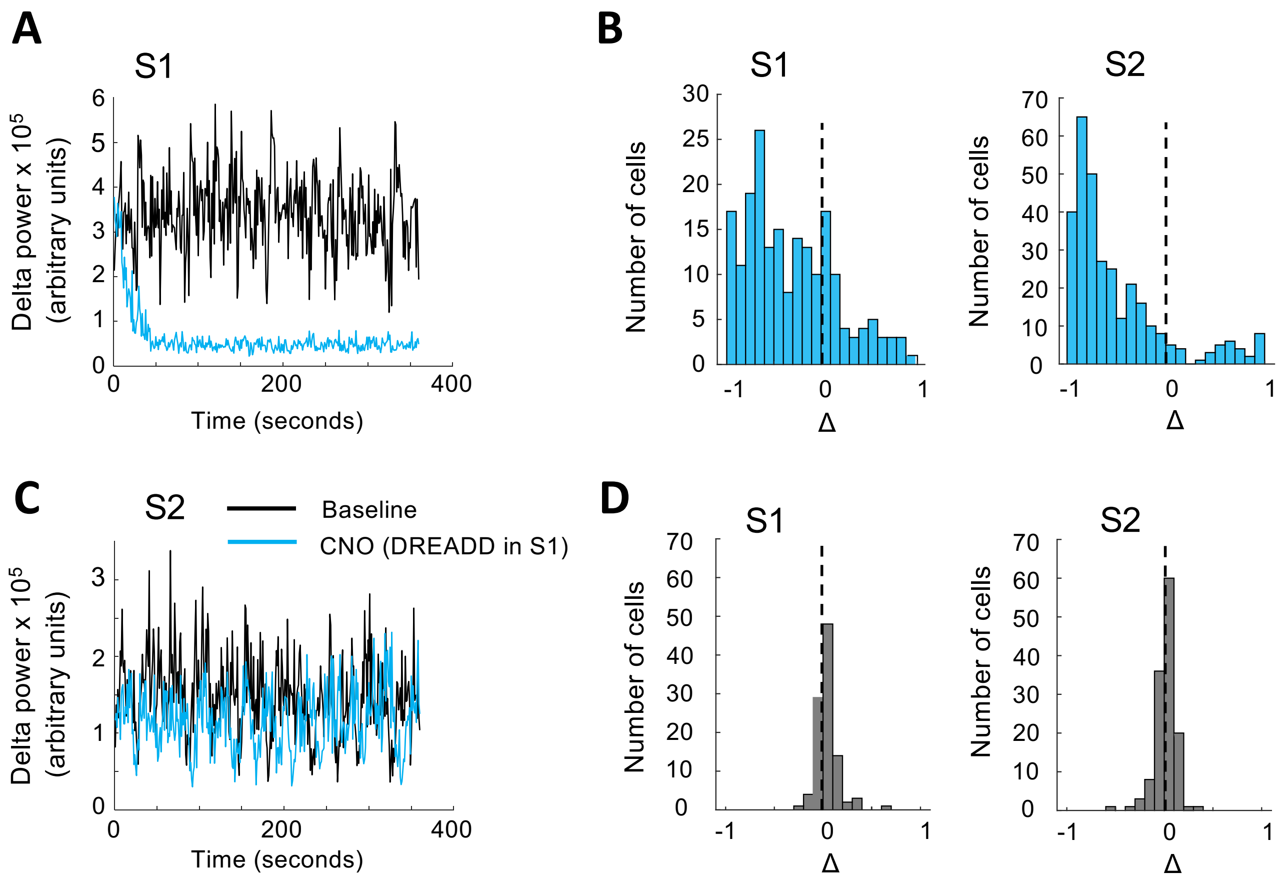


**Supplementary Figure 3.** The effect of DREADD activation on network activity in anaesthetised mouse S1 and S2 cortex. Excitatory hM3Dq is expressed in PV neurones and a mixture of excitatory and inhibitory neurones are recorded using Neuropixel probes. **A:** Activation of S1 expression of DREADDs with CNO causes a rapid decrease in delta-wave activity (0.5-4Hz) shown in blue. Control period activity is shown in black. **B:** Left: Individual neurones discriminated in S1 during CNO administration for S1 DREADD expression. Axis Δ depicts firing change (CNO-baseline)/(CNO+baseline). Zero represents no change. Right: Individual neurones discriminated in S2 during administration of CNO for S2 DREADD expression. In both cases, a large proportion of the cells spontaneous activity is reduced or abolished. A minority of cells increase activity and these are highly likely to be the PV neurones expressing DREADDs. **C:** During S1 DREADD activation, delta activity is reduced slightly in S2. **D:** CNO administration in either S1 (left) or S2 (right) does not lead to a change in firing rate in the absence of DREADDs.

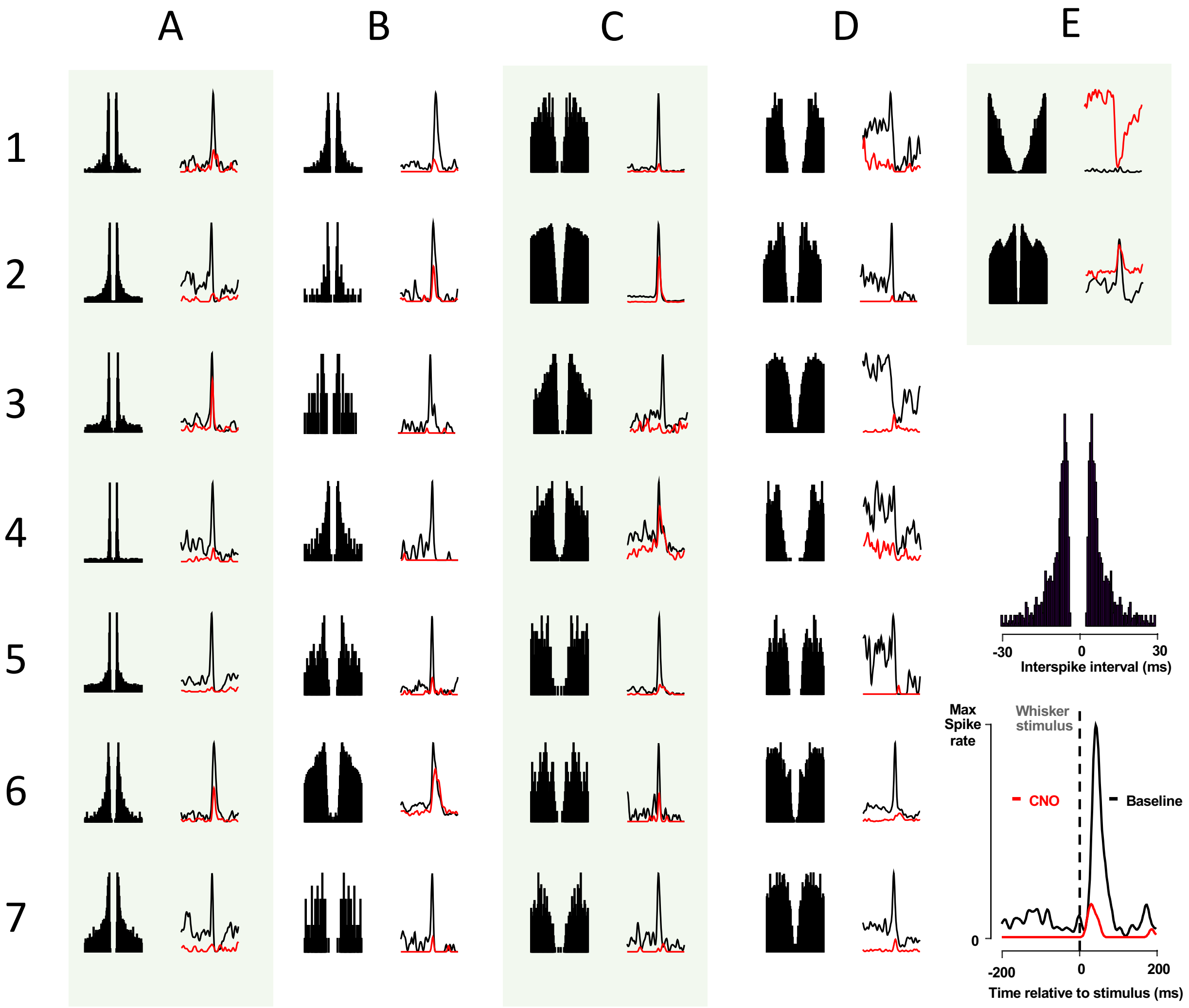


**Supplementary Figure 4. E**ffect of DREADDs on sensory responses in S1. **A-E** (1-7): Examples of the variety of effect of DREADD (red trace) on control firing rate (black). Examples are roughly grouped by inter-interval autocorrelogram (left of each pair). A1-A7 and B1-2 show bursting cells where 6/9 are strongly inhibited. B2-7 neurones with longer intra-burst intervals 4/5 of which are strongly inhibited. Some cells showed high spontaneous activity which was inhibited by DREADD activation (D1-4) three of which had an inhibitory response to stimulation. Two cells are shown that increased their firing rate E1,2 with DREADD activation and are presumably PV neurones. One cell shows an inhibitory response (E1) and the other retained its excitatory response to stimulation at a lower signal to noise ratio E2.

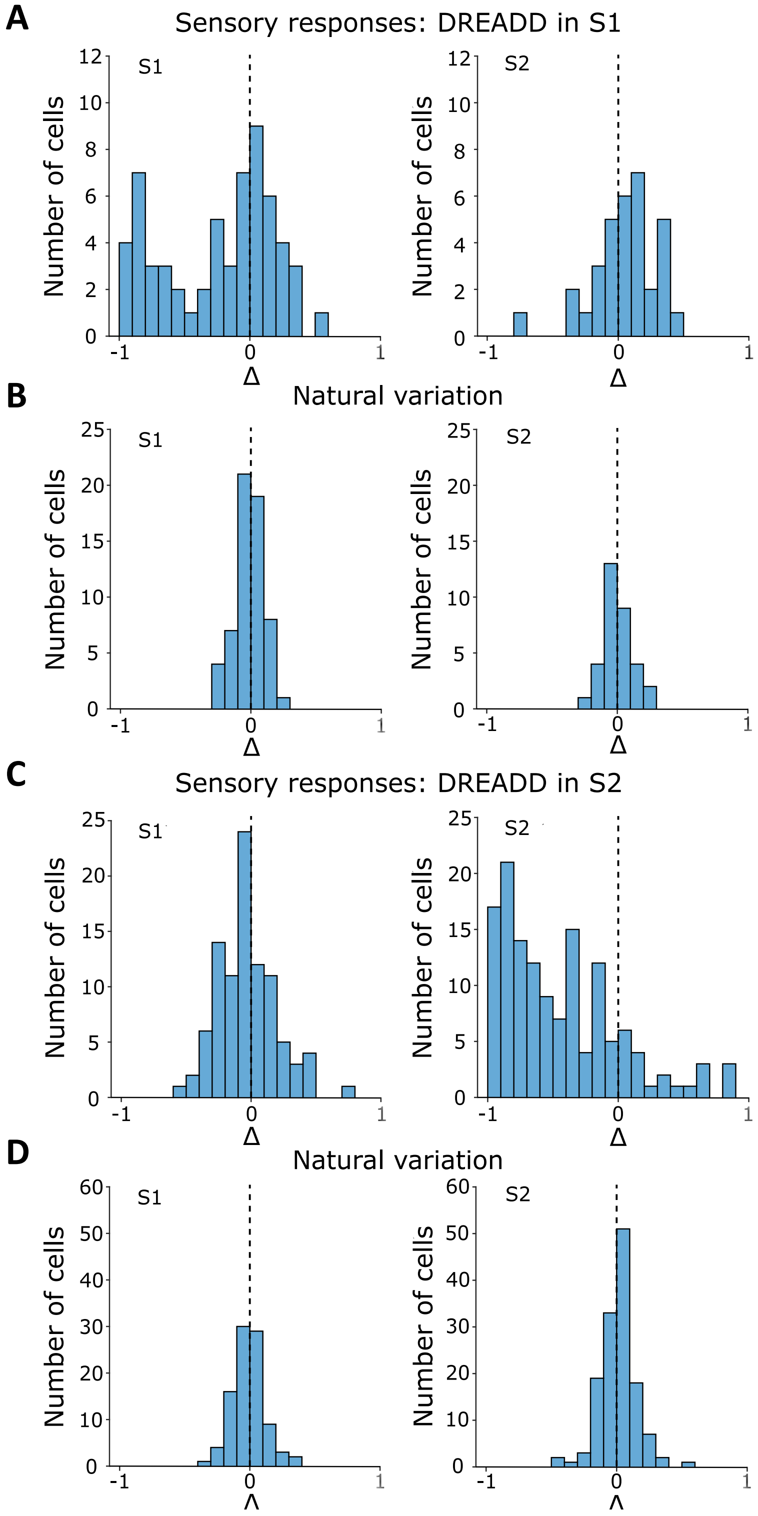


**Supplementary Figure 5.** Effect of DREADDs on sensory responses of just the excitatory cells in S1 and S2. **A:** Effect of DREADD activation in S1 on S1 and S2 neurones. Note that S1 neurones are inhibited (negative Δ values). A few S2 neurones increase their sensory response. **B:** The natural variability in sensory responses are plotted for the same period of time as shown in A, for the baseline period prior to administering CNO. **C:** Effect of DREADD activation in S2 for S1 and S2 neurones. Activation of DREADDs in S2 has a large inhibitory effect on their responses. A few S1 neurones can be seen to decrease their response levels. **D:** Natural variability in sensory responses during the baseline period.

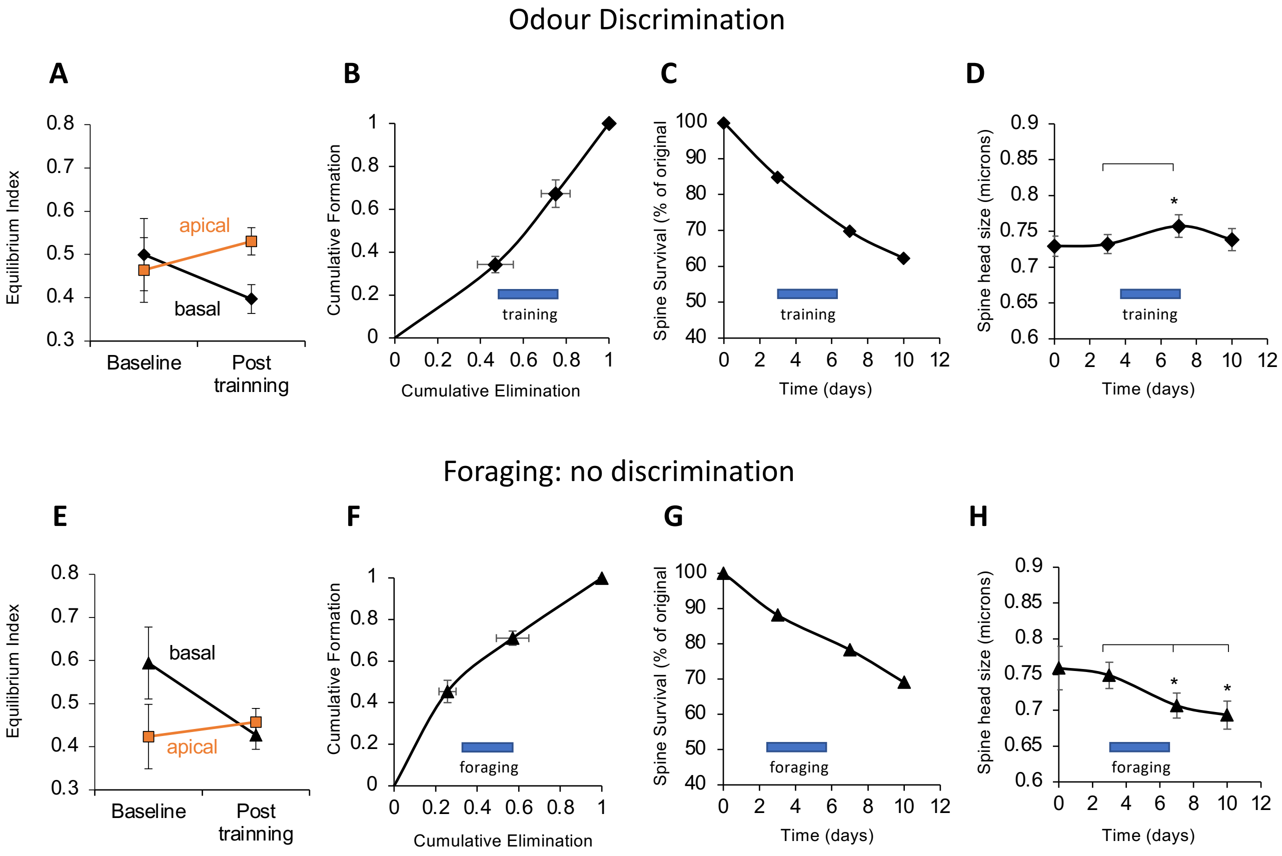


**Supplementary Figure 6.** Two control experiments for the effect of the texture discrimination procedure on spine dynamics in S1. **A-D** spine dynamics for odour discrimination (top) **E-H** Foraging in the absence of discrimination (bottom). **A:** Odour discrimination did not cause a significant change in the equilibrium index (formation/(formation+elimination)) for apical dendrites (orange) or basal dendrites (black) in S1. **B:** The normalised cumulative formation vs cumulative elimination plot is not significantly different from a straight line for basal dendrites. **C:** The rate of spine loss on basal dendrites does not decrease during odour training (blue bar). **D:** Pre-existing spine size does increase slightly during training, but then reverts to control levels. **E:** Foraging in two identical bowls both of which were bated did not cause a significant change in the equilibrium index for apical dendrites (orange) or basal dendrites (black) in S1. **F:** The normalised cumulative formation vs cumulative elimination plot appears to tilt toward greater elimination during foraging (blue bar), but is not significantly different from a straight line for basal dendrites. **G:** The rate of spine loss on basal dendrites does not decrease during foraging (blue bar). **H:** Pre-existing spine size decreases significantly during foraging and remains low even 3 days later. This opposite to the effect seen with texture discrimination.

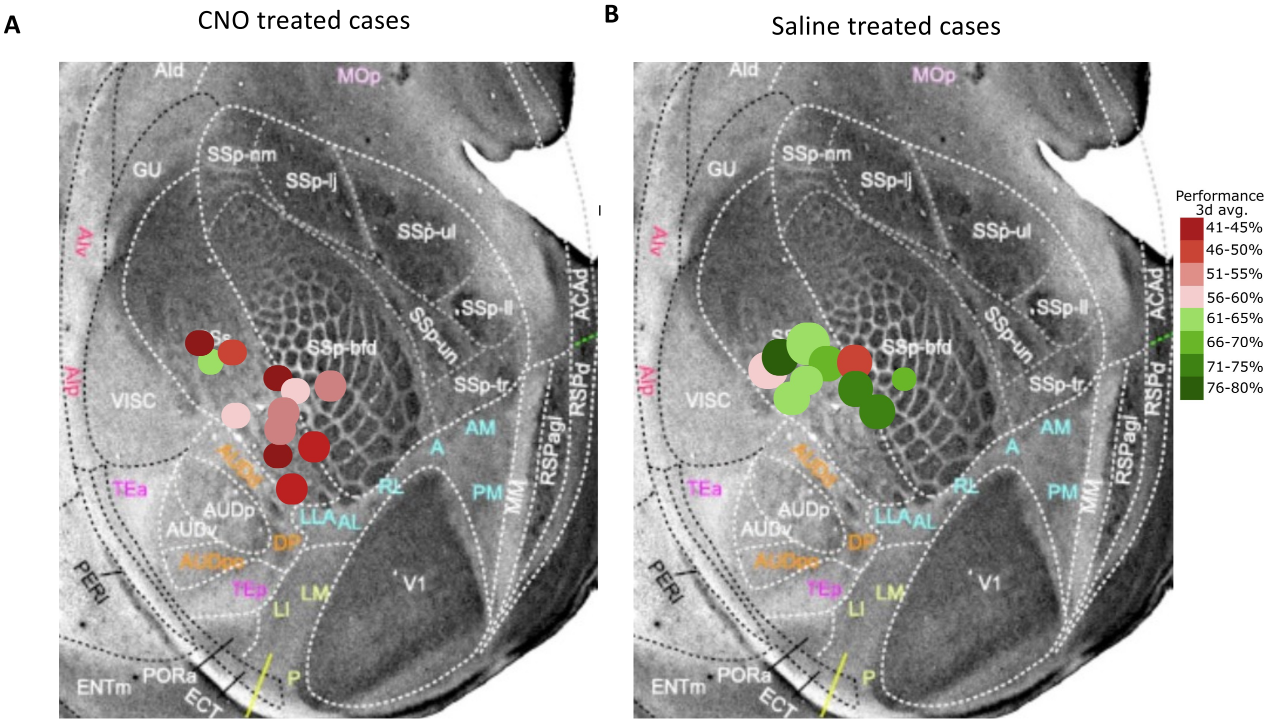


**Supplementary Figure 7.** Location of DREADD expression on a standard map of the flattened mouse brain. The circles are centred on the locus of expression and the colour code indicates the learning performance in the texture discrimination assay (Green = learned, Red = did not learn). **A:** CNO treated cases. Only one mouse learned (green circle in anterior S2). **B:** Saline treated cases. Only two mice did not learn (red circle, pale pink circle). Key on the right shows the percentage of correct trials over the three days (72 trials). Background map of the brain adapted from Gamanut et al. (2018).

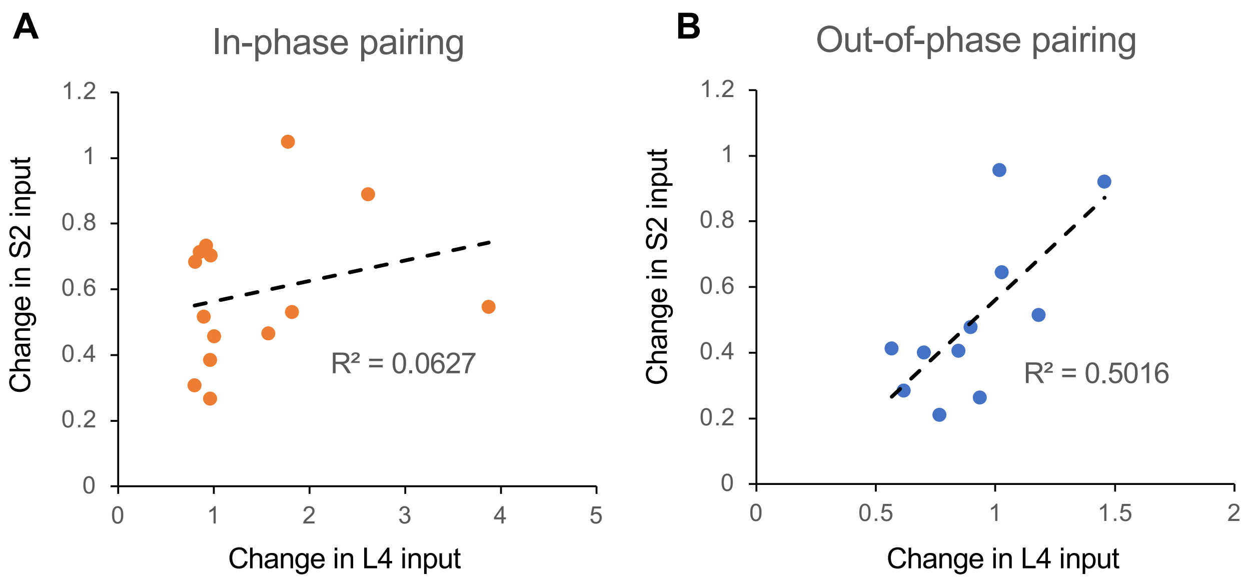


**Supplementary Figure 8:** Degree of correlation between LTD at S1→S2 synapses and change in EPSP amplitude at layer 4→layer2/3 synapses for in-phase and out-of-phase pairing of stimuli. **A:** In phase pairing produces LTD in the S2 input to apical dendrites of L2/3 cells, which is not correlated with the change in L4 input to the same cells' basal dendrites. **B:** Out-of-phase pairing produces LTD in the S2 input as with A, but the amount of LTD is correlated with the size of LTD shown by the L4 inputs to the same cells.
